# Next generation cytogenetics: genome-imaging enables comprehensive structural variant detection for 100 constitutional chromosomal aberrations in 85 samples

**DOI:** 10.1101/2020.07.15.205245

**Authors:** Tuomo Mantere, Kornelia Neveling, Céline Pebrel-Richard, Marion Benoist, Guillaume van der Zande, Ellen Kater-Baats, Imane Baatout, Ronald van Beek, Tony Yammine, Michiel Oorsprong, Daniel Olde-Weghuis, Wed Majdali, Susan Vermeulen, Marc Pauper, Aziza Lebbar, Marian Stevens-Kroef, Damien Sanlaville, Dominique Smeets, Jean Michel Dupont, Alexander Hoischen, Caroline Schluth-Bolard, Laïla El Khattabi

**Affiliations:** Department of Human Genetics, Radboud University Medical Center, Nijmegen, The Netherlands; Radboud Institute of Medical Life Sciences, Radboud University Medical Center, Nijmegen, The Netherlands; Laboratory of Cancer Genetics and Tumor Biology, Cancer and Translational Medicine Research Unit and Biocenter Oulu, University of Oulu, Oulu, Finland; Radboud Institute of Health Sciences, Radboud University Medical Center, Nijmegen, The Netherlands; Department of Cytogenetics, University hospital of Clermont-Ferrand, France; Department of Medical Genetics, Cochin Hospital, APHP.Centre, University of Paris, Paris, France; lnstitut Neuromyogène, CNRS UMR 5310, INSERM U1217, Lyon 1 University, Lyon, France; Unit of Medical Genetics, Saint-Joseph university, Beyrouth, Lebanon; Department of Genetics, Hospices Civils de Lyon, Bron, France, Institut Neuromyogène, CNRS UMR 5310, INSERM U1217, Lyon 1 university, Lyon, France; Cochin Institute, INSERM U1016, Paris, France; Department of Internal Medicine and Radboud Center for Infectious Diseases (RCI), Radboud University Medical Center, Nijmegen, The Netherlands

## Abstract

Chromosomal aberrations and structural variations are a major cause of human genetic diseases. Their detection in clinical routine still relies on standard cytogenetics, karyotyping and CNV-microarrays, in spite of the low resolution of the first one and the inability to detect neither balanced SVs nor to provide the genomic localization or the orientation of duplicated segments, of the latter. We here investigated the clinical utility of high resolution optical mapping by genome imaging for patients carrying known chromosomal aberrations in a context of constitutional conditions.

For 85 samples, ultra-high molecular weight gDNA was isolated either from blood or cultured cells. After labeling, DNA was processed and imaged on the Saphyr instrument (Bionano Genomics). A *de novo* genome assembly was performed followed by SV and CNV calling and annotation. Results were compared to known aberrations from standard-of-care tests (karyotype, FISH and/or CNV-microarray).

In total, we analyzed 100 chromosomal aberrations including 7 aneuploidies, 35 translocations, 6 inversions, 2 insertions, 39 copy number variations (20 deletions and 19 duplications), 6 isochromosomes, 1 ring chromosome and 4 complex rearrangements. High resolution optical mapping reached 100% concordance compared to standard assays for all aberrations with non-centromeric breakpoints.

Our study demonstrates the ability of high resolution optical mapping to detect almost all types of chromosomal aberrations within the spectrum of karyotype, FISH and CNV-microarray. These results highlight its potential to replace these techniques, and provide a cost-effective and easy-to-use technique that would allow for comprehensive detection of chromosomal aberrations.

## Introduction

Structural variants (SV) play an important role in human diversity and diseases. The emergence of cytogenetic tools, starting with karyotyping followed by fluorescence *in situ* hybridization (FISH) and CNV-microarrays, allowed for their detection and thereby significantly contributed to the discovery of disease causing genes.^1-3^ However, these tools show significant limitations as karyotyping has a very low resolution, estimated at 5-10 Mb on average. Additionally, CNV-microarrays are not able to detect mosaicism lower than 5-20% or balanced chromosomal aberrations, and do not provide information on the location of the structural variation, e.g. mapping of insertions is impossible.

Despite their drawbacks, karyotyping and CNV-microarrays still represent major tools in the routine genetic investigation of constitutional and somatic diseases, since chromosomal aberrations are major causes of e.g. reproductive disorders, recurrent miscarriages, congenital malformations or (neuro-)developmental disorders. Karyotyping is thereby indicated for diseases where numerical and structural balanced aberrations are highly represented, such as in reproductive disorders where (sex)chromosomal aneuploidies and large structural aberration including balanced rearrangements are frequently present.^4-7^ CNV-microaray is recommended as first-tier test for developmental disorders (DD) with or without multiple congenital anomalies (MCA),^8^ as it enables the diagnosis of sub-chromosomal copy number variations (CNV) including clinically relevant microdeletions/microduplications.^1; 2; 9^ In DD/MCA, the diagnostic rate rose from less than 5% with karyotyping^10; 11^ to 15 to 20% with CNV-microarray leading to the replacement of the former analysis by the later as a first-tier test.^8; 12^

The recent breakthrough in sequencing technologies raised great interest in complementing or replacing cytogenetic tools for an all-in-one genetic test allowing for the detection of both nucleotide variants and structural variants.^13-15^ Moreover, short read sequencing became reasonably inexpensive and is versatile in terms of protocols (gene panel, whole exome sequencing (WES) and whole genome sequencing (WGS)). Yet, the detection of structural variants remains challenging because of (i) the relatively limited read length and (ii) the repetitive nature of sequences at some structural variation breakpoints. Although many improvements regarding technical aspects and data analysis pipelines have been achieved, genome sequencing is still not able to comprehensively and cost-effectively detect balanced structural anomalies, impeding its wide implementation in clinical cytogenetic laboratories. Moreover, the most comprehensive analysis of SVs in WGS data requires the use of multiple tools, as established e.g. by the 1000 genomes project SV consortium. ^16^ Hence, a real-time analysis with fast turnaround time is not yet feasible for each and every laboratory. It is expected that long-read whole genome sequencing (LR-WGS) will dramatically improve the ability to identify SVs in individual genomes,^17^ and examples have shown this utility for individual research cases.^18; 19^ However, the routine use of long-read sequencing as a diagnostic tool requires several improvements.

To this end, a tool complementary to sequencing that may truly replace standard cytogenetics may offer great additional value. Optical mapping by genome imaging consists of imaging very long linear single DNA molecules (median size >250 kb) that have been labeled at specific sites. Since its first description, ^20^ this formerly tedious technique has been updated by Bionano Genomics. They combined microfluidics, high-resolution microscopy and automated image analysis to allow for high-throughput whole genome imaging and its *de novo* assembly.^21; 22^ Historically, such maps have been used as a scaffold to guide the assembly of NGS contigs to build reference genomes of several plant and animal species.^23-25^ More recently, methods dedicated to the detection of SVs in humans have been developed. Data analysis thereby includes two distinct pipelines: a CNV pipeline that allows for the detection of large unbalanced aberrations based on normalized molecule coverage, and an SV pipeline that compares the labeling patterns between the constructed genome maps of the studied sample and a given reference. The latter allows for the genome-wide detection of SVs as small as few hundred base pairs, including insertions, deletions, duplications as well as inversions and translocations.

Optical mapping using Bionano^®^ recently proved to allow for efficient detection of a wide range of chromosomal anomalies in leukemia.^26^ It has also been used to detect germline SVs in individual research cases^27; 28^ or individuals from the 1000 genomes consortium^16^ and to unravel population specific SVs.^29^

The aim of the current study was to benchmark Bionano Genomics’ optical mapping technology against standard-of-care cytogenetic tools (karyotype, FISH and/or CNV-microarray). To do so, we analyzed a wide range of simple and challenging chromosomal aberrations, which had been previously characterized by standard approaches, in samples from patients with a broad range of clinical indications.

## Subjects and methods

### Patient selection and sample collection

This multicenter study involved a total of 85 samples from four genetic academic centers from the Netherlands (Radboud University Medical Center, RUMC) and France (Cochin hospital in Paris, Hospices Civils in Lyon and the university hospital of Clermont-Ferrand). Patients were referred to one of the inclusion centers for developmental or reproductive diseases. Recommended chromosomal investigations were performed according to the indications. Karyotyping was performed in case of reproductive disorders or family history of balanced chromosomal anomaly. CNV-microarray, and karyotyping for some samples, was performed in case of developmental disorders. In some cases, additional investigations including fluorescence *in situ* hybridization (FISH) were performed to characterize an identified anomaly.

Cases for which (i) a chromosomal anomaly was identified by karyotyping, CNV-microarray or FISH, and (ii) for which there was enough residual blood (EDTA or heparin) or cultured cells available after routine testing, were included. Samples were anonymized or informed consent is available, respectively. Blood samples for high molecular weight DNA extraction were stored at −20°C for a maximum of one month and at −80°C for longer term storage. In addition, several cases with known aberrations had material other than blood available as a residual material from routine testing. This included 8 amniotic fluid cell lines, 4 chorionic villi cell lines and 8 lymphoblastoid cell lines, which were all generated from primary cultures according to standard diagnostic procedures.

### Karyotyping

Karyotyping was performed according to previously described standard protocols.^30^ Chromosomal abnormalities were described according to the International System for Human Cytogenetic Nomenclature (ISCN, 2016).

### Fluorescence in situ hybridization

Fluorescence *in situ* hybridization (FISH) was performed on standard chromosome slides according to the manufacturer’s instructions (Vysis, Abbott, USA), or using isolated BAC-clones as FISH-probes following standard procedures.

### CN V-microarray

CNV-microarray was performed using the Agilent SurePrint G3 ISCA v2 CGH 8×60K or SurePrint G3 Human CGH Microarray 4×180K (Agilent Technologies, Santa Clara, CA, USA), or the Affymetrix Cytoscan ™ HD Array (Thermo Fisher Scientific, Waltham, USA). Genome coordinates were provided according to hg19/GRCh37 human reference genome.

### Ultra-high molecular weight DNA isolation, DNA labeling and data collection for optical mapping

For each patient, ultra-high molecular weight (UHMW) DNA was isolated from 400 μL of whole peripheral blood (EDTA or Heparin) or 1-1.5 million cultured cells (lymphoblastoid cells, amnion cells or chorionic villi cells), using the SP Blood & Cell Culture DNA Isolation Kit and according to manufacturers’ instructions (Bionano Genomics^®^, San Diego, CA, USA). Briefly, cells were treated with LBB lysis buffer to release genomic DNA (gDNA) which was bound to a nanobind disk, washed and eluted in the provided elution buffer.

UHMW DNA molecules were labeled using the DLS (Direct Label and Stain) DNA Labeling Kit (Bionano Genomics^®^, San Diego, CA, USA). Direct Label Enzyme (DLE-1) and DL-green fluorophores were used to label 750 ng of gDNA. After a wash-out of the DL-Green fluorophores excess, DNA backbone was counterstained overnight before quantitation and visualization on a Saphyr^®^ instrument.

Labeled UHMW gDNA was loaded on a Saphyr chip^®^ for linearization and imaging on the Saphyr instrument (Bionano Genomics, San Diego USA) (Supplementary Figure 1).

### De novo assembly and structural variant calling

The *de novo* assembly and Variant Annotation Pipeline were executed with Bionano Solve software v3.4 or v.3.5. Results were analyzed through two distinct pipelines: a CNV pipeline that allows for the detection of large unbalanced aberrations based on normalized molecule coverage, and an SV pipeline that compares the labeling patterns between the constructed sample genome maps and a reference genome map. Reporting and direct visualization of structural variants were performed using Bionano Access software v1.4.3 or v.1.5.1. The following filtering thresholds were applied: confidence values for insertion/deletion=0, inversion=0.01, duplications= −1, translocation=0 and CNV=0.99. SV calls were compared to an optical mapping dataset of 204 human control samples (provided by Bionano Genomics) to filter out common SVs and potential artifacts (both technical and reference-genome related) (Supplementary Figure 1).

### Data analysis and comparisons

All optical mapping results were analyzed genome-wide (Supplementary Table 1) for all samples irrespective of the patient’s chromosomal status. We subsequently compared SVs and CNVs detected by optical mapping to the ones previously identified by standard-of-care techniques (karyotype and/or CNV-microarray).

## Results

### Population description

All 85 samples included in this study were previously analyzed by karyotyping, FISH and/or CNV-microarray according to the reason for referral and the respective international recommendations (Figure 1A and B, Supplementary Table 1). Reasons for referral included developmental delay including autism spectrum disorders or intellectual disability, associated or not with congenital malformations (49 patients, 57.6%), reproductive disorders (15 patients, 17.6%), familial history of chromosomal aberration (12 patients, 14.1%), and abnormal prenatal test results (9 patients, 10.6%). These samples exhibited a total of 100 chromosomal aberrations with 11 different types of aberrations from the previous standard diagnostics tests, summarized in Figure 1C. Additionally, nine known aberrations in this cohort were beyond the scope of this study due to breakpoints in the (peri-)centromeric regions of any chromosome or p-arm of acrocentric chromosomes, and were therefore excluded from further analyses.

**Figure 1.**
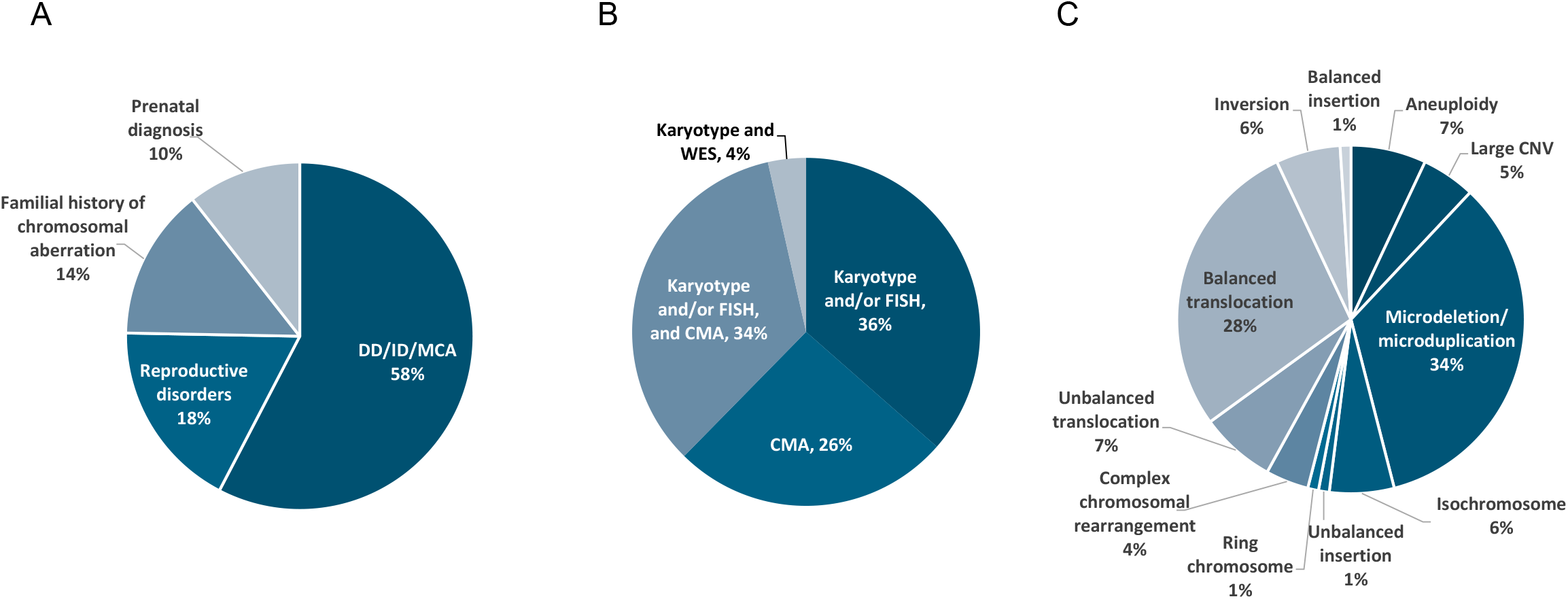
Description of the study population (n=85). A) Main reason for referral. B) Distribution of the different cytogenetic and molecular tests used for diagnosis. C) Distribution of chromosomal aberrations as assessed by standard of care genetic investigations.

### Results of Optical Mapping with Bionano Genome Imaging

Bionano genome imaging generated on average 655 Gbp of data per sample (853 Gbp for samples processed in Nijmegen, aiming at ~200X genome coverage, and 463 Gbp for samples processed in France, aiming at ≥80X genome coverage per sample, respectively). The average N50 molecule length (> 150 Kbp) was 267 Kbp and label density was 15.1 labels/100 Kbp. This resulted in an average map rate of 76.8% and an effective coverage of 152x (192x for Radboud samples, 114x for French samples) (Supplementary Table 2).

Structural variant calling identified on average 5,758 (+/− 344) SVs per sample, of which the vast majority corresponded to insertions and deletions (with an average of 4,127 (+/− 239) and 1,549 (+/− 108) respectively). Filtering out events which were present in a database comprising of 204 population control samples resulted in an average of 80 (+/−65) rare SVs per sample, of which 41 (+/− 28) were overlapping with genes (Figure 2, Supplementary Table 3). Besides SV detection, CNV detection was performed using a separate coverage-depth based algorithm that is included in the *de novo* assembly and variant calling pipeline.^31; 32^ This analysis resulted in an average of 1 gain and 10 losses per sample without applying any size threshold cut-offs (Supplementary Figure 2). Of note, CNV calls are often segmented into multiple calls, hence the true number of CNVs is expected to be lower.

**Figure 2:**
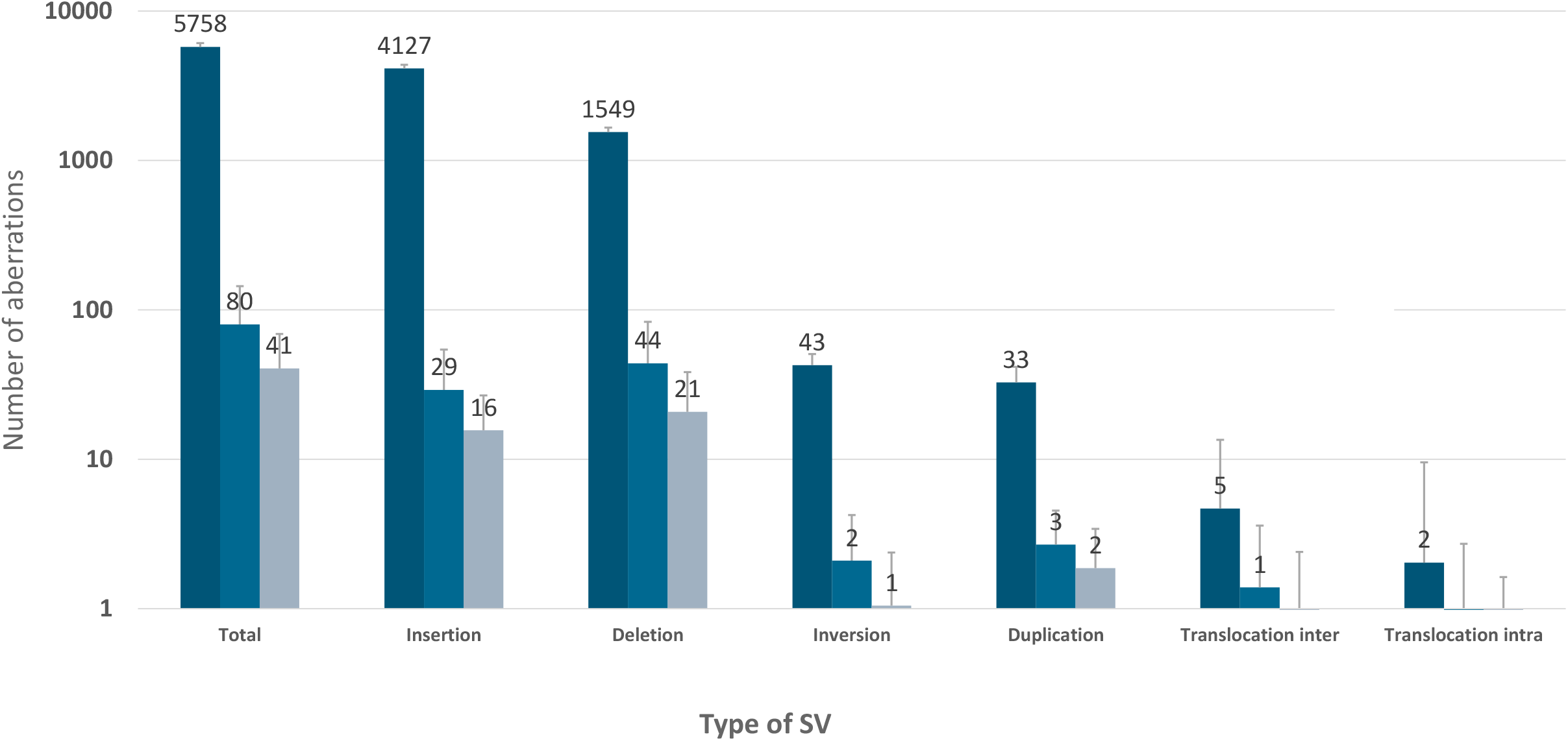
SV detection and filtering. Average number of SVs detected per sample, given per type of SV (total, insertion, deletion, inversion, duplication, interchromosomal translocation and intrachromosomal translocation). Dark blue: all variants, median blue: rare variants only (not found in control database including 204 samples), light blue: rare variants that overlap with genes.

### Detection of diagnostically reported aberrations with genome imaging

All diagnostically reported aberrations in our study cohort were detected correctly either by the SV or the CNV calling, with several aberrations being identified by both algorithms, reaching a 100% concordance for optical mapping with the previous diagnostic test results (Supplementary Table 1). For five samples however, filter settings needed to be adapted in order to detect the expected aberrations (see Supplementary Table 1). Adaptation included setting the confidence value for CNVs to 0 (3 samples) and turning off the SV DLE-1 mask filter (2 samples).

The 100 identified aberrations included 7 aneuploidies, 20 deletions and 19 duplications, 35 translocations, 6 inversions, 2 insertions, 6 isochromosomes and 1 ring chromosome (Figure 1C). In addition, four of our patients showed complex chromosomal rearrangements, defined as cases where aberrations involve three or more chromosomes or when at least four SVs are detected on the same chromosome. For graphical representation of different types of chromosomal aberrations see Figures 3 and 4.

**Figure 3:**
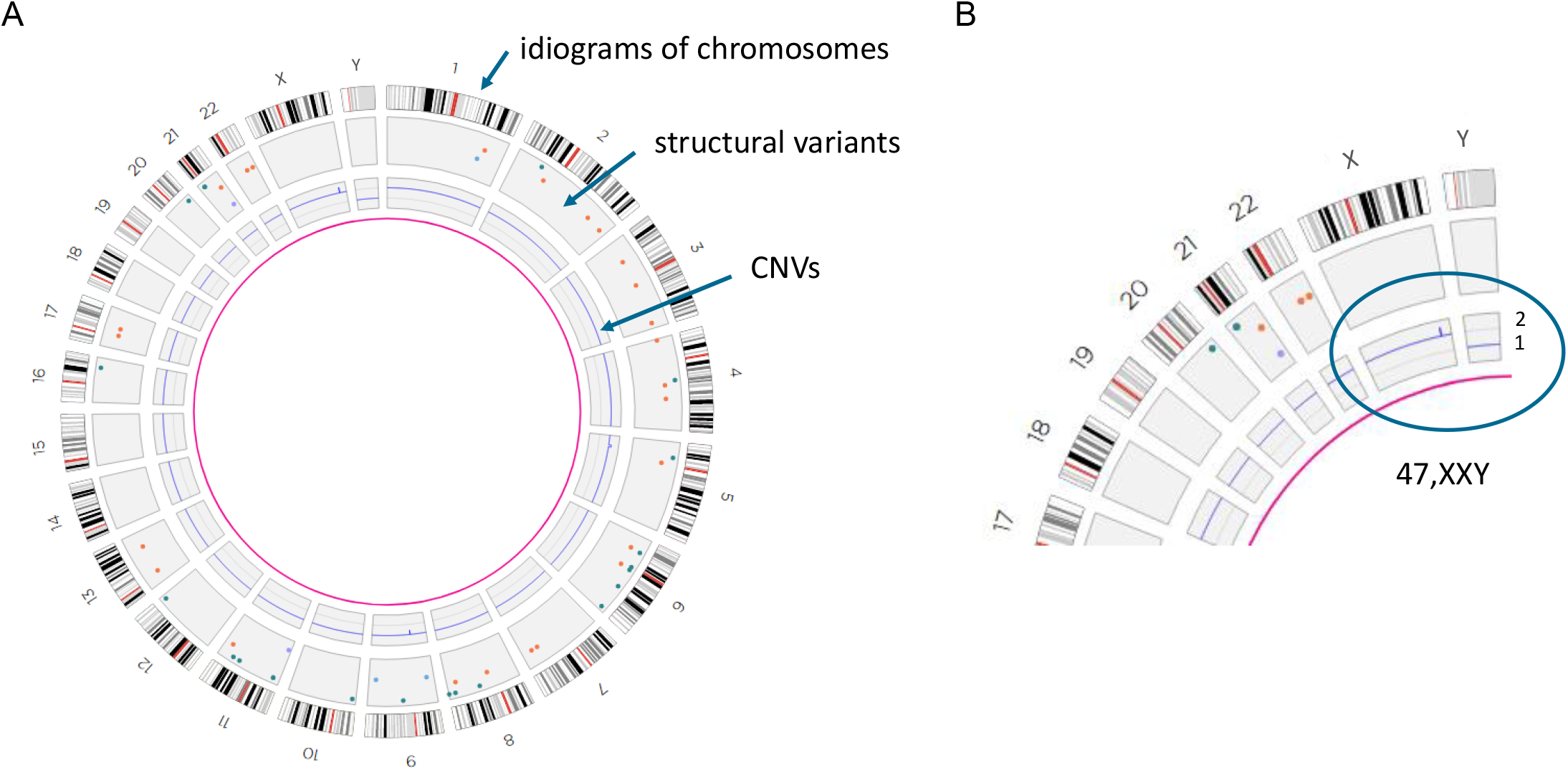
Visual representation of optical mapping data A) Genome-wide circos plot showing all 24 chromosomes in a circular way. For each chromosome, the idiograms are shown at the outside of the circosplot, with ideogram-style chromosomal banding and the centromeres in red. Different colored dots in the boxes underneath represent different called SVs. The blue line in the box underneath represents the CNV profile, with each peak representing a CNV call. B) Part of a circos plot, showing the sex chromosomes. The blue CNV line shows two copies of chromosome X, as for autosomes, and one copy of chromosome Y consistent with a sex chromosome aneuploidy (47,XXY, resulting in Klinefelter syndrome).

**Figure 4:**
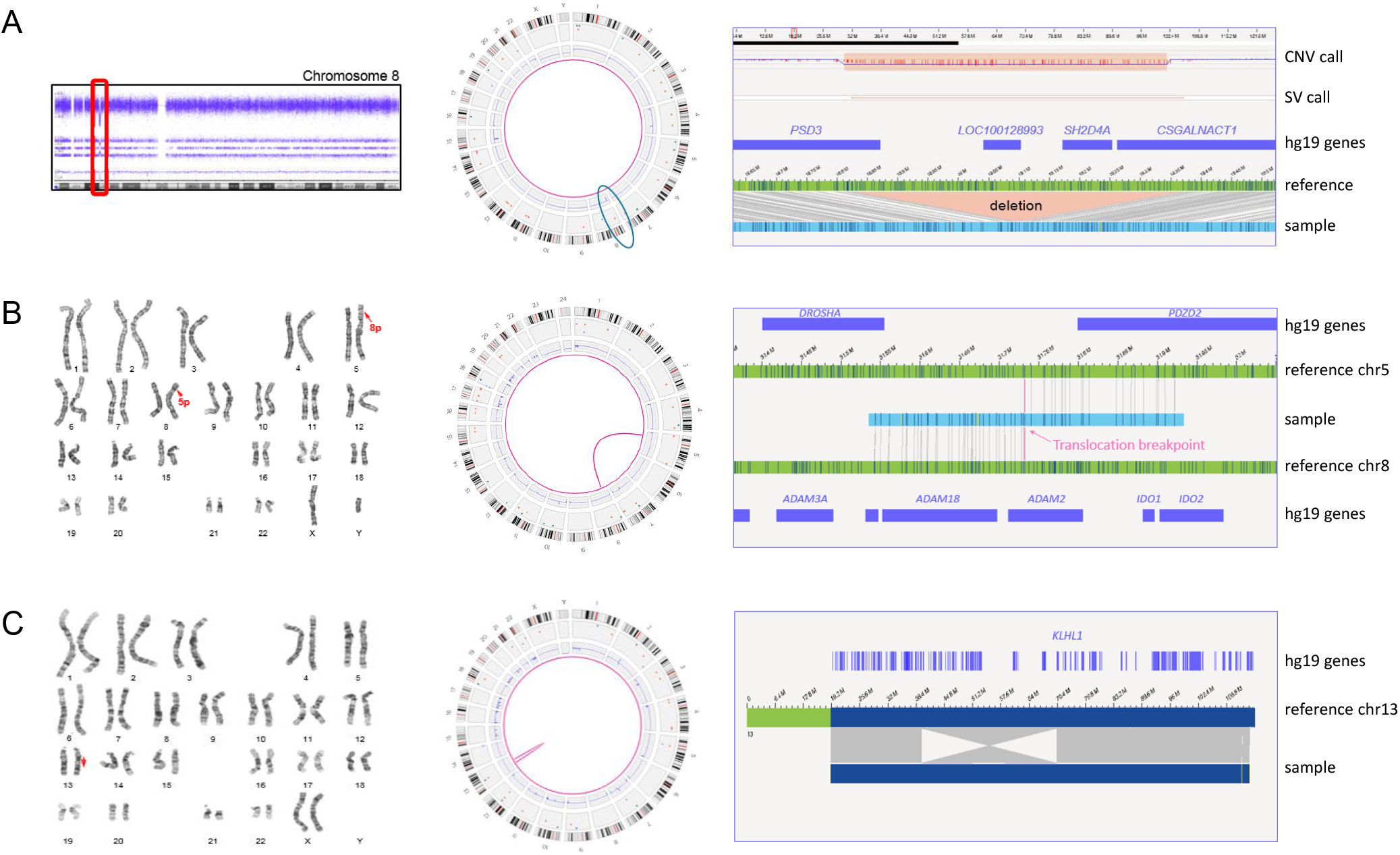
Representation of different chromosomal aberrations. A) Sample 1. Left: CNV microarray data showing an 8p22p21.3(18825888_19364764) deletion. Middle: Genome-wide circos plot showing all chromosomes. The deletion is detected by the CNV and the SV pipeline (blue circle). Right: genome map with reference, showing the deletion and affected genes (hg19). B) Sample 18. Left: karyogram showing a 46,XY,t(5;8)(p13.1;p11.2) karyotype. Middle: circos plots with a pink line connecting chromosomes 5 and 8, representing the translocation. Right: genome map, of which the left part maps to chromosome 8 and the right part to chromosome 5. C) Sample 15. Left: karyogram showing an inversion on chromosome 13 (red arrow). Middle: circos plot showing the inversion as an intrachromosomal translocation. Right: genome map that is partly inverted when compared to the reference. One of the breakpoints is interrupting the gene *KLHL1*.

### Aneuploidies, partial aneuploidies and large CNVs

Our study cohort included 7 full aneuploidy samples, including 3x XXY, 2x monosomy X, 1x trisomy 14 and 1x trisomy 21 (the two latter ones were detected in prenatal samples and were mediated by Robertsonian translocations). In addition, 4 mosaic monosomy X samples were included (Supplementary Table 1). All aneuploidies of the autosomes were called correctly with the used algorithms, whereas the aneuploidies of the sex chromosomes had to be manually inferred from the visualized data of the CNV plot (Figure 3). This manual inference is no longer required with the recent Bionano Solve v3.5. In addition to whole chromosome aneuploidies, five large CNVs ranging in size between 6.6 and 14 Mb, and 7 large aberrations corresponding to derivative chromosomes from unbalanced translocations detected by karyotyping, were included and detected correctly.

### Isochromosome

Six of our samples contained isochromosomes. Four of those were iso-dicentric Y-chromosomes, one sample contained an isodicentric chromosome 15, and another had an isodicentric chromosome X. The four isodicentric Y-chromosomes all showed a similar genome map pattern (Figure 5). Whereas all four have normal coverage at the p-arm and a small part of the q-arm (until q11.221), there is no coverage at q11.222 to q11.23 and at the end of q12. The largest part of q12 had no coverage in none of the samples (including controls), as this part of the chromosome represents a gap in the reference genome (hg19, N-base gap). Interestingly, whereas samples 27, 57, and 79 have a nearly identical coverage pattern, only sample 55 shows a slightly different breakpoint, with a part of q11.222 still being covered. While the CNV or coverage pattern undoubtedly allows to decide about the presence of isochromosomes in all samples, it should be noted that centromeres itself cannot be detected, hence the distinction between dicentric vs. monocentric status may remain uncertain in some cases. Of note, the isochromosomes of chromosomes 15 and X presents are remarkable: For chromosome 15, the fractional copy numbers of the affected regions differed, and were called as 3 and 4. The isodicentric chromosome X was present in low mosaic state (see Supplementary Figure 3).

**Figure 5:**
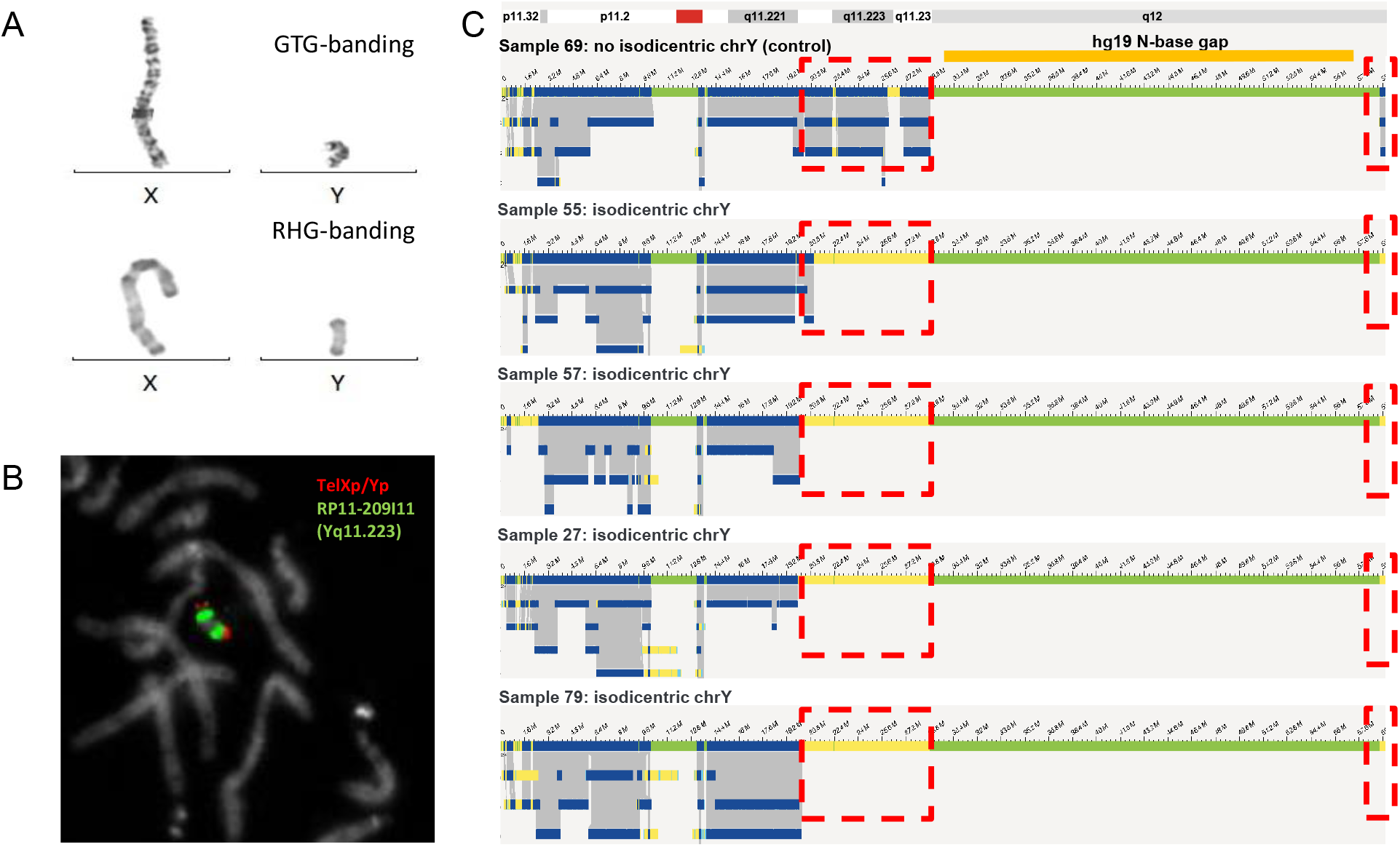
Isodicentric Y-chromosomes show specific bionano assembly map patterns. A) GTG-and RHG banding of X- and Y-chromosomes of sample 57. B) FISH for sample 57 using probes TelXp/Yp and RP11-209I11 (Yq11.223). C) Bionano genome maps of Y-chromosomes of samples 69 (no isodicentric chrY), 55, 57, 27 and 79. Dotted red boxes indicate where isodicentric Y-chromosomes have no coverage when compared to non-isodicentric Y chromosomes (the top genome map).

### Ring chromosome

One of the samples analyzed contained a mosaic ring chromosome X, as previously detected by karyotyping (Figure 6). The karyotype reported was 45,X[14]/46,X,r(X)(p11.21;q21.1)[21]. The patient presented with growth retardation and development delay. Following genome imaging, an intrachromosomal translocation on chromosome X was detected, connecting positions chrX: g.57,009,891 (p11.21) and chrX:g.78,599,384 (q21.1), confirming and refining the positions previously detected by karyotyping. The fractional copy number of 1.6 for this region, compared to 1.0 for the rest of this chromosome confirmed the mosaic state of this ring chromosome.

**Figure 6:**
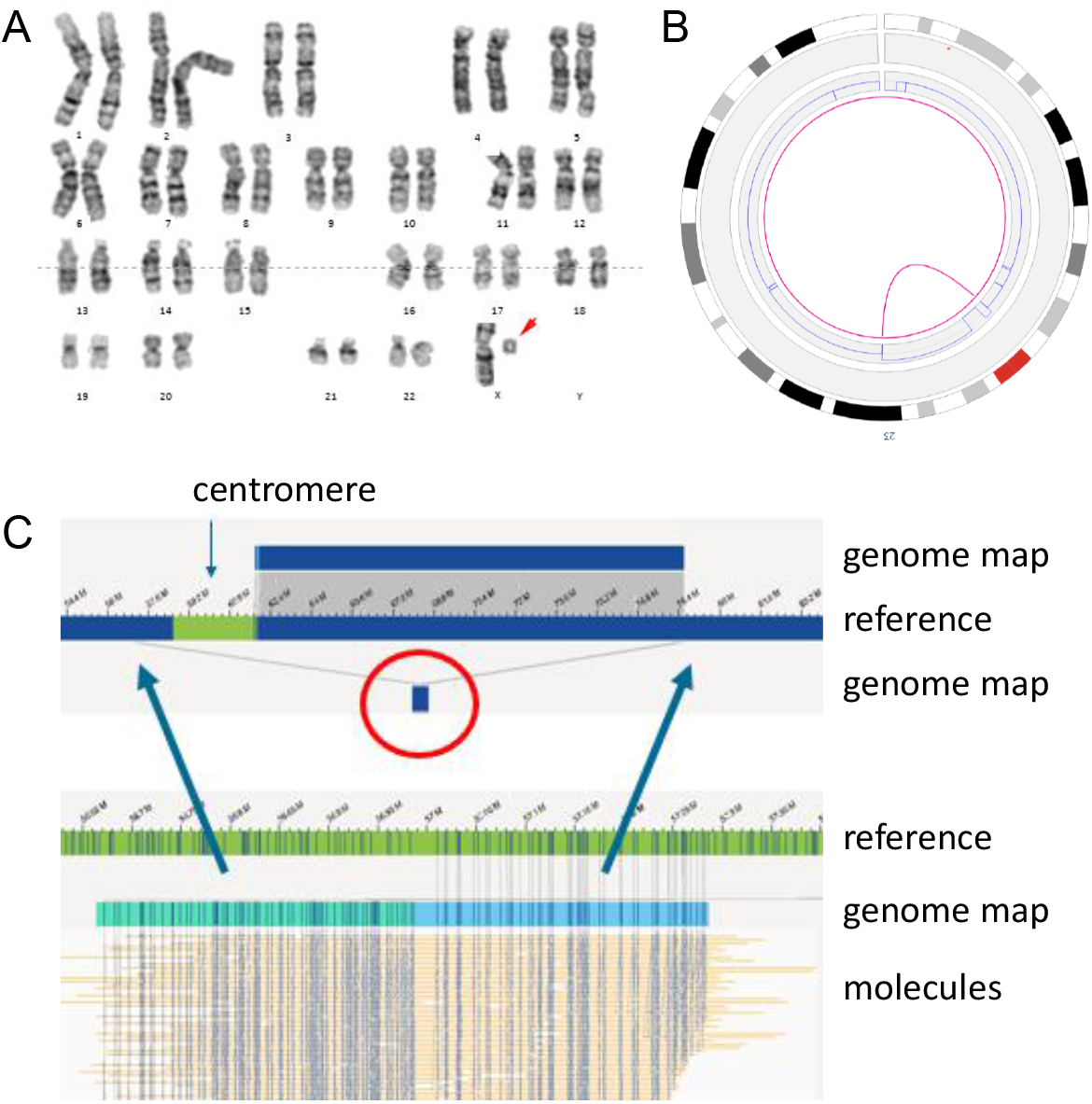
Small X ring chromosome. A) Karyogram of sample 39. The red arrow is pointing towards the small X ring chromosome. B) Circos-plot (of chromosome X only) of sample 39. The pink line in the center of the circosplot is indicating the presence of the ring chromosome (called as an intrachromosomal translocation). C) Different genome maps (dark blue bars on top and below the reference) indicating the presence of the ring chromosome. The individual molecules for the genome map below the reference (highlighted by a red circle) are shown at the bottom of this figure. The left part of these molecules (light green bar) map to a region upstream of the centromere, whereas the right part of the same molecules (light blue bar) map to a region downstream of the centromere.

### Translocations and inversions

Thirty-five of the investigated samples carried previously identified balanced (n=28) and unbalanced (n=7) translocations, which were all detected by genome imaging. As expected, unbalanced translocations were detectable by both structural variant calling and CNV calling, whereas balanced translocations and inversions were only detected by SV calling (Figure 4B, C). Traditionally, balanced translocations can be detected via karyotyping but not via CNV-microarray. Genome imaging is able to refine translocation breakpoints for such cases. Accordingly, several balanced translocations and inversions were shown to likely disrupt protein-coding genes, including the well described *SETBP1* (MIM: 611060), *KANSL1* (MIM: 612452), *DYRK1A* (MIM: 600855), and *PIGU* (MIM: 608528) genes, with the latter two being disrupted by the same translocation (Figure 7). The breakpoints for *KANSL1* (sample 49) had previously been validated using FISH and whole-genome sequencing (WGS),^33^ whereas the others are newly uncovered and still need to be confirmed. In all cases, the patient’s phenotype matches the expected phenotype for the dominant diseases associated with the respective genes. Detection of breakpoints with optical mapping is much more accurate than karyotyping. For the few breakpoints for which WGS data were available for comparison,^33^ the breakpoint accuracy was within five kb (Supplementary Figure 4).

**Figure 7.**
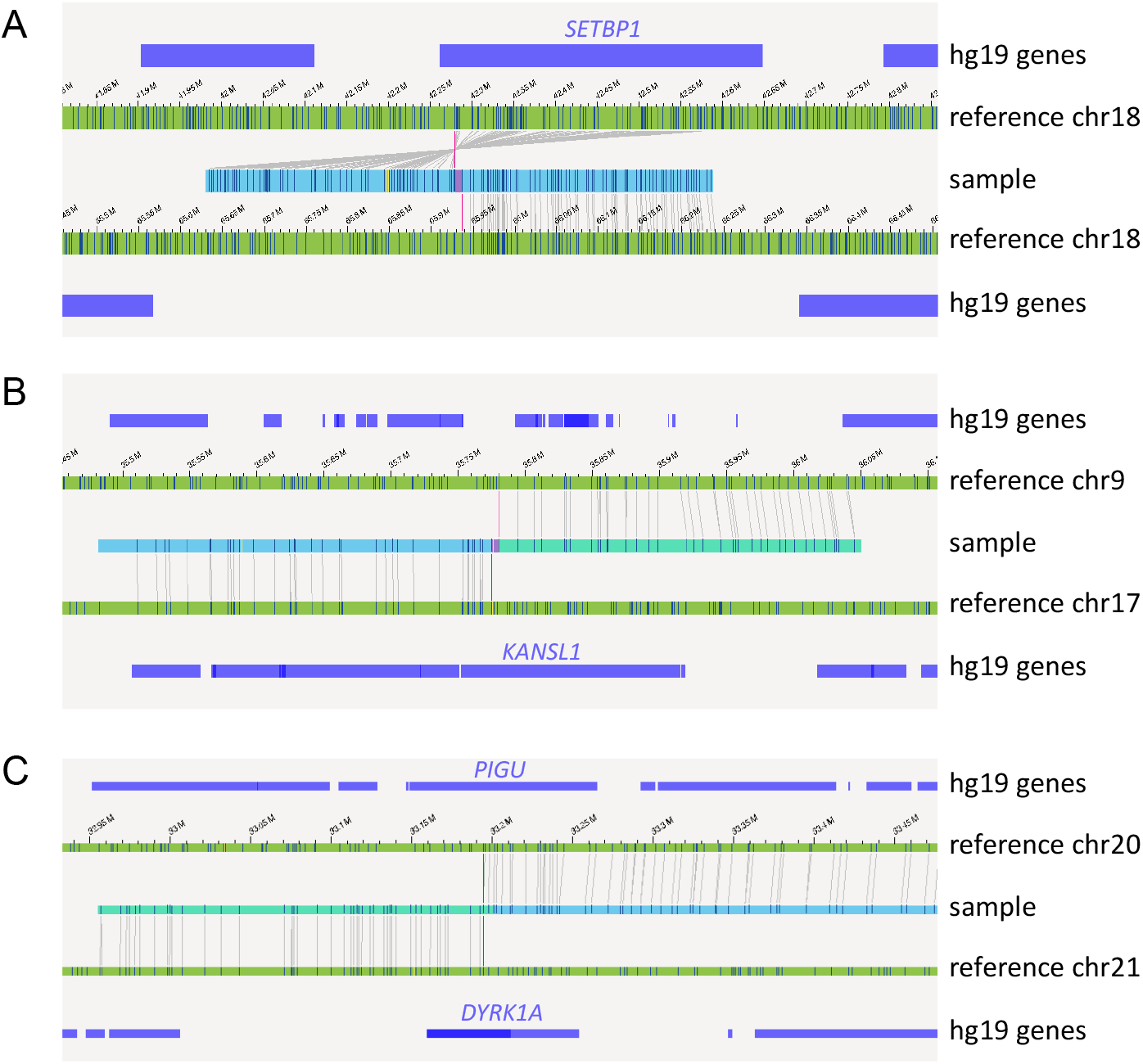
Examples of inversions and translocations interrupting well known disease causing genes. A) Inversion inv(18q)(q22.1q12.3), disrupting the *SETBP1* gene in sample 47. B) Translocation t(9;17)(p13.3;q21.31), interrupting the gene *KANSL1* in sample 49. C) Translocation t(20;21)(q11.22;q22.13), interrupting the genes *DYRK1A* and *PIGU* in sample 54.

### Microdeletions and -duplication

In addition to large chromosomal aberrations (aneuploidies, large CNVs and translocations), our cohort included 34 microdeletion/-duplications (<5 Mb). These microdeletion/duplications ranged in size from 34 Kbp (sample 84) to 4.2 Mbp (sample 44), and included some of the well-known microdeletion/duplication syndromes such as DiGeorge syndrome (22q11.2 deletion syndrome, OMIM: 188400), Williams-Beuren syndrome (deletion 7q11.23, OMIM: 194050), Charcot-Marie-Tooth syndrome type 1A (CMT1A, duplication 17p12, OMIM: 118220) and 1q21.1 susceptibility locus for Thrombocytopenia-Absent Radius (TAR) syndrome (OMIM: 274000). Although the presence of segmental duplications (SegDups) for several of these microdeletions/duplications often leads to breaking of the genome maps, all microdeletions/duplications were correctly called by either the SV or CNV algorithms or both, although most events were called by the CNV tool. SeqDups are often mediating recurrent CNVs, for example the 22q11.2 microdeletion-causing DiGeorge syndrome (Figure 8). Depending on size and structure of these SeqDups, the SVs were occasionally disrupted, or falsely filtered out due to high population frequencies of partially overlapping SVs (Supplementary Table 1). We expect that the analysis of individual molecules of sizes up to 2Mb shall allow full assembly maps even for those regions but additional software improvements may be required.

**Figure 8.**
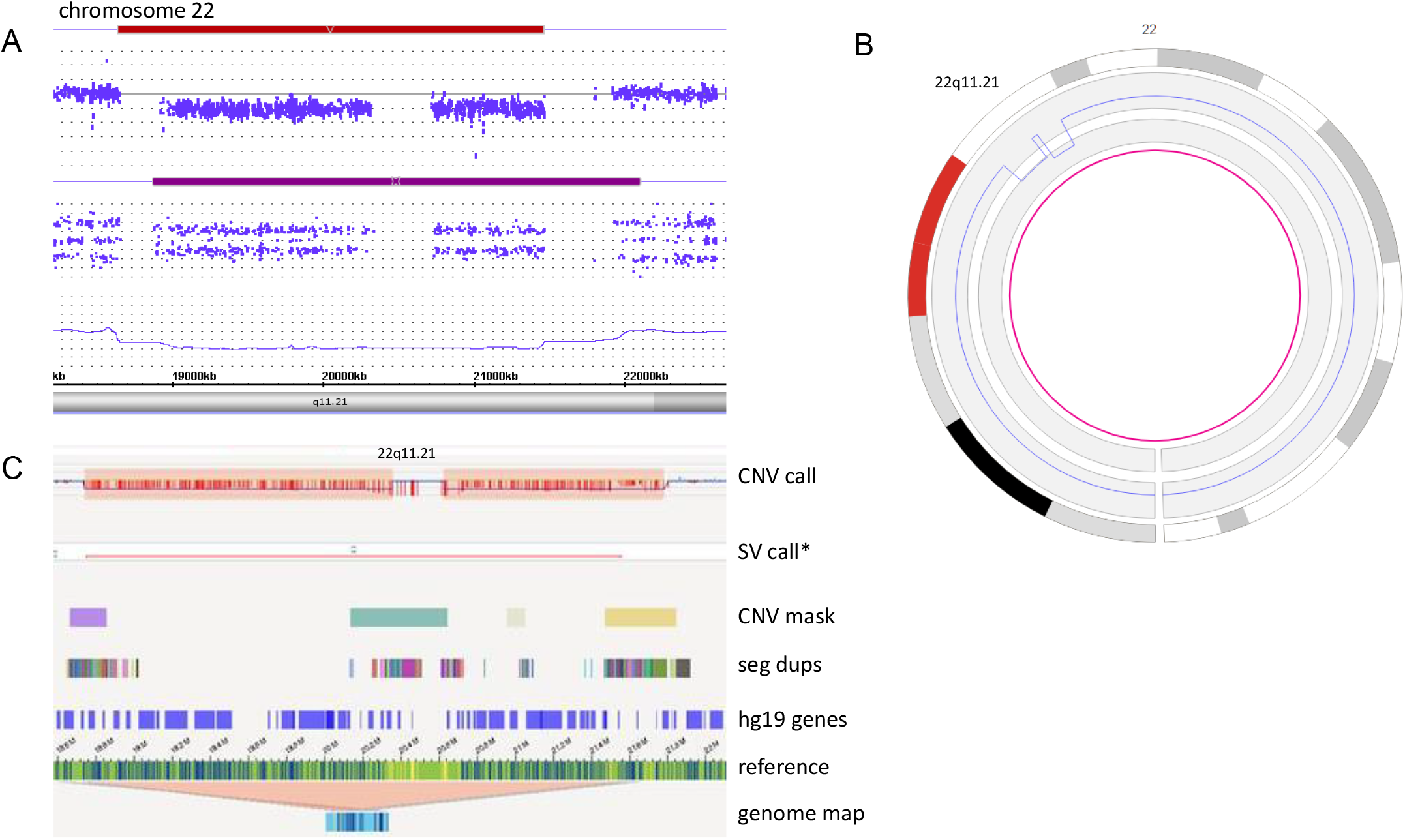
Example of a typical 22q11.2 microdeletion syndrome, in sample 2 (VCF, Di-George syndrome). A) Affymetrix CytoScan HD array, showing the 22q11.21(18308819_18519921)x1 deletion. B) Bionano circos plot, showing an aberrant CNV profile on 22.q11.21. C) Bionano genome maps of chr22q11.21, showing the CNV calls (on top), a segmental duplication bed file (below), the chromosome 22 reference genome map (again below) and the different genome maps (at the bottom). The deletion is clearly called by the CNV calls, and is surrounded by segmental duplications. *SV not called with standard filters, but recovered when % of the Bionano control sample overlap was set to 80%.

### Complex cases

Finally, four of the samples included in this study presented with complex rearrangements (28, 52, 55, 66), four of which are samples of patients with developmental delay and/or intellectual disability (Supplementary Table 1). For example, karyotype of Sample 28 (Figure 9) showed a translocation t(3;6)(q1?2;p2?2), a derivative chromosome 4 (?der(4)(:p1?2->q1?2:)) and a derivative chromosome 5 (der(5)(4pter->4p1?2::4q1?2->4q34.2::5p14.2->5qter)) in different clones. CNV-microarray showed losses on 4q34 (4q34.2q34.3(176587929_190957474)x1 dn) and 5p15 (5p15.33p14.2(113577_24449849)x1 dn). Following optical mapping, the translocation t(3;6)(q1?2;p2?2) was identified as t(3;6)(q13.12;p24.3). In addition, a translocation t(4;5)(q34.2;p14.2), a loss of 4q34.2q34.3 and a loss of 5p15.33p14.2 were detected, concordant with previous results. In the same sample, genome imaging also revealed putative additional translocations t(3;4)(q13.11;q12), t(3;4)(q13.11;p11) t(4;6)(q12;p22.3) and an inversion inv(13)(q31.2;q33.3) (Figure 9).

**Figure 9:**
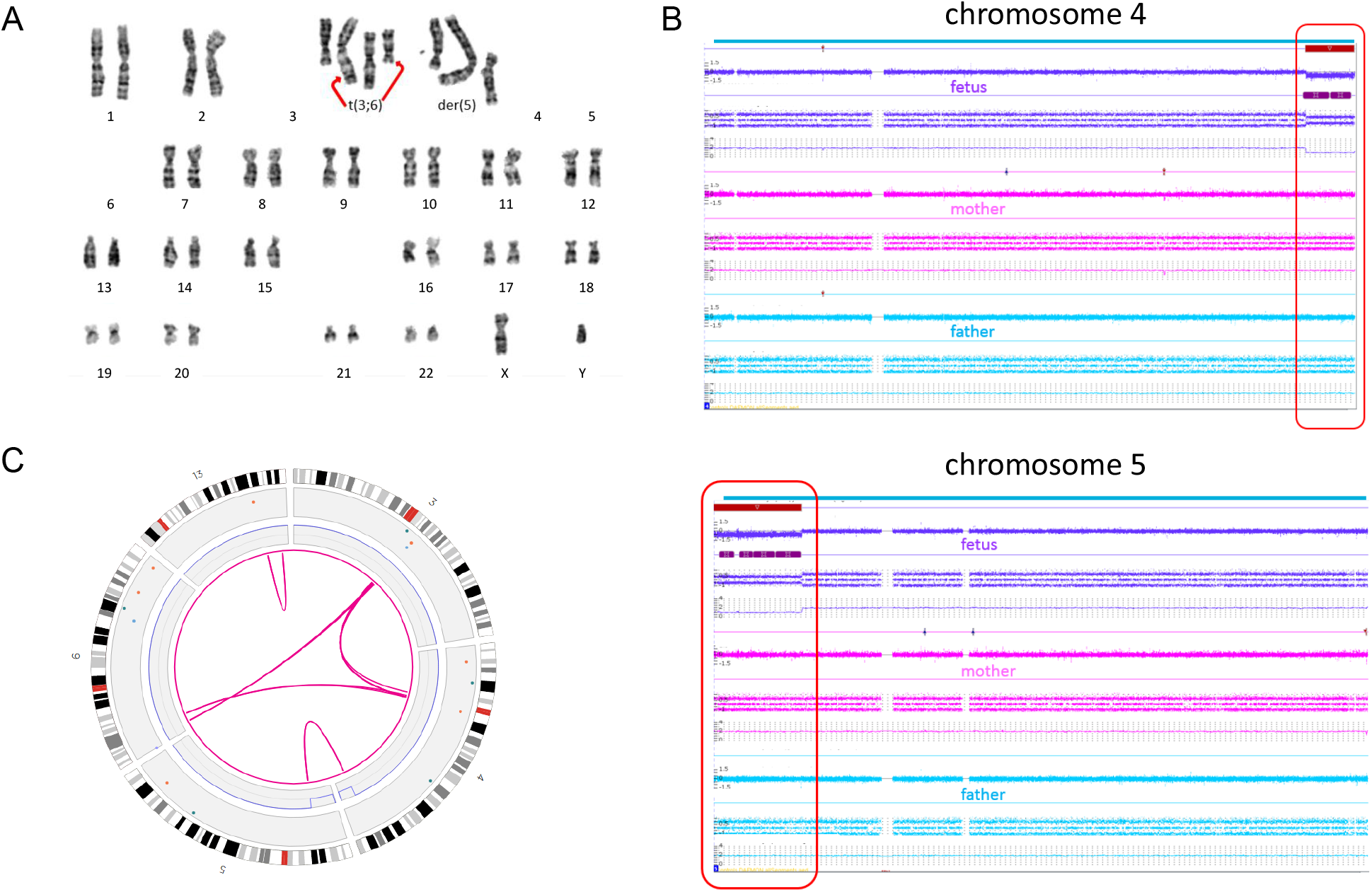
Complex sample 28. A) Karyogram showing the translocation t(3;6) and the derivative chromosomes 4 and 5. B) CNV microarray data showing two *de novo* deletions on chromosome 4 and 5. C) Bionano circos plot of chromosomes 3, 4, 5, 6 and 13. The Bionano data confirm all previous data and show the presence of additional translocations t(3;4) and t(4;6) plus an inversion on chromosome 13.

Another sample (66) showed a 3-way translocation t(3,13,5)(p11.1;p12;p14) after karyotyping and four losses on chromosome 3 (3p14.1(65238298_68667113)x1,3p13(70127345_73724765)x1, 3p12.1(83784489_85467284)x1,3q11.2(97180779_97270083)x1) following CNV-microarray (Figure 10). Optical mapping confirmed these aberrations, but unraveled additional complex rearrangements on chromosome 3, leading to the identification of a chromoanagenesis. For all residual samples with complex rearrangements, see Supplementary Table 1 and Supplementary Figures 5 and 6.

**Figure 10:**
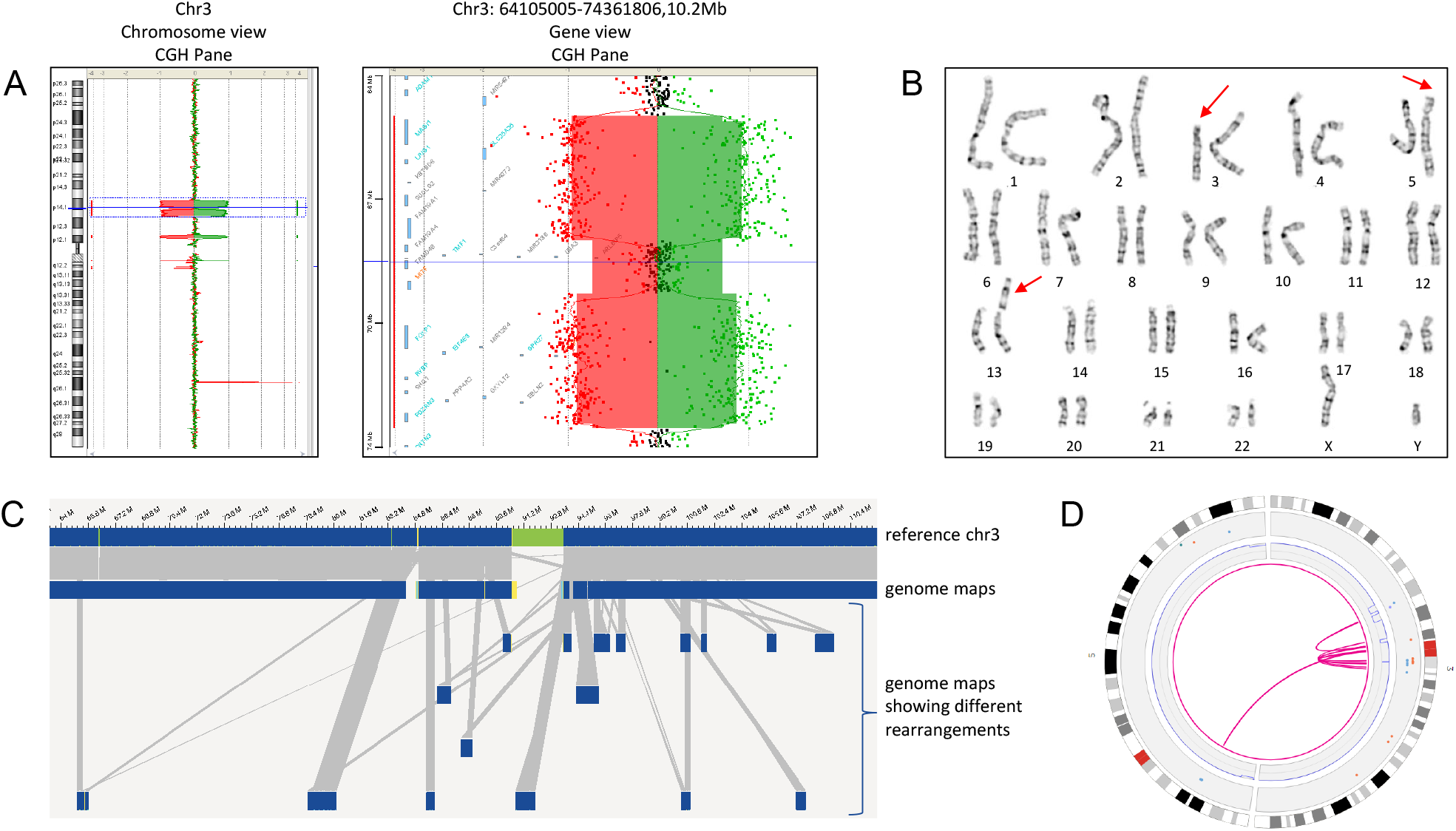
Complex sample 66. A) CGH Panes of sample 66, showing the whole chromosome 3 (left) and chr3:64105005-74361806 (right). B) Karyogram of sample 66: 46,XY,t(3;13;5)(p11.1;p12;p14). C) Genome maps of chromosome 3 of sample 66, showing multiple rearrangements on chromosome 3. D) Circos plot of chromosomes 3 and 5, showing multiple intrachromosomal translocations on chromosome 3 and a translocation between chromosomes 3 and 5.

Taken together, optical mapping allowed the correct unravelling of complex karyotypes, which previously required the combination of karyotyping, FISH and CNV-microarrays, by combining the detection of translocations and imbalances (CNVs, gains and loss of genetic material) including balanced and unbalanced events in one assay and at an unprecedented resolution.

## Discussion

Chromosomal aberrations and SVs are frequently involved in many genetic diseases including developmental disorders, congenital malformations, intellectual deficiency, reproductive disorders as well as cancer. Hence, their accurate detection is critical to achieving a complete genetic investigation but limitations of the current standard-of-care genetic analyses (karyotyping, FISH, CNV-microarray and NGS) preclude any comprehensive characterization without combining multiple approaches.^16; 34^ Indeed, to date, not a single technology offers a full resolution of chromosomal aberrations in all samples. The traditional karyotyping is still performed as the first-tier test in case of reproductive disorders in spite of its poor diagnostic rate (overall less than 10%), likely due to its very low resolution. Moreover, its quality is unpredictable since it varies between samples and laboratories, it depends on the availability of viable cells and relies on the expertise of the technician and the cytogeneticist which is decreasing over the years because of lack of training. Hence, there is need for a more robust, high-resolution and automatable method. On the other hand, CNV-microarray represents one such robust routine tool, that has allowed an improved diagnostic yield e.g. to approximately 15% for neurodevelopmental disorders,^8^ but it lacks the ability to detect balanced aberrations such as translocations or inversions or to decipher the orientation of duplicated or inserted segments, and resolution remains restricted to a few kilobases. Sequencing based assays for SV detection are constantly improving.^35; 36^ This includes improved CNV calling from exome or genome sequencing, however most comprehensive detection requires a combination of analysis tools.^37; 40^ Moreover, it requires the local implementation of bioinformatic pipelines that have not yet undergone a large scale clinical validation. In addition to technical and computational hurdles, SV detection by sequencing based technologies becomes difficult when breakpoints localize within repetitive sequences which is frequently observed since many SVs are caused by the non-allelic homologous recombination of repeats in the first place. It is expected that long-read sequencing may enable near perfect genomes one day, but so far technologies, analyses as well as throughput and prices do not allow its routine clinical use.^17^

In this manuscript, we have shown that genome imaging is capable of comprehensively and easily detecting all classes of chromosomal aberrations and may complement or replace current cytogenetic technologies. Our cohort was composed of a large panel of different tissues, aberrations and indications representative of what can be encountered in clinical routine. We demonstrated that optical mapping allows for the detection of balanced as well as unbalanced rearrangements at sizes ranging from few kilobases to several megabases or even entire chromosome aneuploidies. The method allows for detection of SVs down to 500 bp, but none of our clinically reported SVs were that small. Copy number variations and aneuploidies were all detected by either the coverage analysis pipeline and/or structural variation analysis. Combining two analysis pipelines, one based on coverage depth and the other one based on the comparison of a *de novo* assembled genome map with a reference map, allows for the most complete detection of all balanced and unbalanced aberrations, as shown by our results. In fact, the first pipeline performs better for large deletions and duplications, and is currently the only tool to detect terminal chromosomal gains/losses or other events that do not create the fusion of unique novel molecules. The second pipeline is more sensitive to small CNVs down to few hundred base pairs, and allows for best breakpoint resolution.

Regarding translocation-, inversion- or insertion-events, we demonstrate that they can all be detected as long as the breakpoints do not lie within large repetitive, unmappable regions such as centromeres, p-arm of acrocentric chromosomes, or heterochromatin stretches. Challenges to map such breakpoints along with the inability of the current software to detect loss of heterozygosity were known prior to this study and such results were expected. In fact, these regions are likely not well represented in the human reference genome,^41^ they cannot be specifically labeled and they are several megabases long, far larger than the longest single molecules that can be obtained with any current technology. Interestingly, in some cases we were able to detect translocations with breakpoints that lie in pericentromeric regions and which were not detected by paired-end whole-genome sequencing (samples 50, 51 and 54; NGS data not shown, manuscript in preparation). This additional detection did solve the molecular diagnosis for patient 54 whose karyotype is 46,XY,t(20;21)(q11.2;q21). Optical mapping showed that this balanced translocation disrupts the *DYRK1A* gene and refined the breakpoint to 21q22.13. This patient displays autism spectrum disorder and microcephaly consistent with a *DYRK1A* haploinsufficiency, which has been shown to be associated with autism spectrum disorder, intellectual disability and microcephaly.^42-44^ Similarly, other cases of gene disruption provided hints into the molecular diagnosis of intellectual disability. For example, patient 47 had an inversion that we showed to likely disrupt the *SETBP1* gene and patient 49 had a t(9;17) that potentially disrupts *KANSL1* as previously identified by WGS.^33^ In both cases, haploinsufficiency of the respective genes is known to lead to clinical syndromes including intellectual disability ^45; 46^ consistent with our patient’s phenotypes.

Optical mapping was also able to detect complex rearrangements including multiple translocations, or even chromoanagenesis. In some cases, optical mapping results suggested a more complex event than expected (28, 47, 52, 66, 70, 74) where the additional SV calls need to be further validated. Furthermore, in four cases of isochromosome Y, the CNV pipeline detected a coverage profile that is very suggestive of an isochromosome (samples 27, 55, 57, 79) similarly to or better than CNV-microarray results, although the Bionano SV pipeline did not call the isochromosome Y. Such isochromosomes with breakpoints in the long arm of chromosome Y are not detectable by sequencing technologies.

As a non-sequencing based technology, single molecule optical mapping overcomes issues due to repetitive regions inaccessible to sequencing. For some aberrations, it even enabled the detection of breakpoints mapping at segmental duplications (Figure 8).^47^

It is not unlikely that at some stage (long read) sequencing approaches may allow fully comprehensive assessment of all SVs and chromosomal aberrations in each personal genome, possibly after *de novo* genome assembly instead of re-sequencing.^41; 48^ Some benefits of optical mapping may prevail: 1.) relative ease of analysis, 2.) relative low costs, 3.) optical mapping can produce 300-1600X genome coverage allowing the reliable detection of rare somatic events, with additional improvements in development.^26^

From a technical point of view, optical mapping using Bionano can best be compared to an ultra-high resolution karyotype (~10,000 times higher resolution than the conventional karyotype) that offers a fast (3-4 days from sample to variant calls) and cost-effective (~$450 list price per genome) alternative to both karyotyping and CNV-microarray. Neither significant data storage capacities nor bioinformatics processing are required. Turnaround time has also been significantly improved in the recent years, as six genomes can be processed in a single run, and instrument price has been reduced ($150k list price). It is also worth noting that the filter settings suggested here result in a small number of events per case (n= 41 rare SVs on average), while detecting all previously known events. This is suggestive for a low false positive rate, although orthogonal validations e.g. by sequencing were beyond the scope of this study. This may be in contrast to NGS-based SV calling: several reports point out the high number of false calls with sequencing based technologies.^49; 50^ Additional clinical analysis filter may include overlap-analysis of SVs with known disease genes or loci. This is a crucial point since in the context of clinical routine, genetic investigation should be time-efficient since the longer the turn-around time the lower the quality of disease management especially in case of reproductive disorders. In addition, our results support the robustness of the technology as our samples were processed in three different facilities. Results were highly similar in terms of quality metrics, number of variant calls and performance stated by the rate of concordance with conventional cytogenetic analyses. Some differences in the number of calls are most likely due to different versions of analysis software used. Clinically relevant results were unchanged.

The technology has the potential to keep improving at both technical and analytical levels. Indeed, a closer examination of the maps or loosening the filters for few samples (34, 42, 50, 76, and 81) led to the identification of initially missed structural variants, supporting the potential of improvement of the software or analysis settings. Other aspects that are being improved to meet cytogeneticist expectations include loss of heterozygosity analysis, ideogram style representation of chromosomal aberrations, e.g. translocations, ISCN nomenclature outputs, and hyperlinks to genome databases. As with comparative genomic hybridization, polyploidies cannot be detected with the current analysis pipeline but haplotype analysis should make this detection possible.

The main focus here was to investigate the concordance i.e. true positive rate for known aberrations as a first step to explore the possibility to replace standard cytogenetic assays by optical mapping. In addition, it is also attractive that this can complement NGS to achieve a better and possibly nearly complete genome analysis. In developmental disorders, optical mapping could complement sequencing approaches to allow for a comprehensive genomic investigation. In reproductive disorders, it could replace karyotyping as the main method complemented by a count of few metaphase spreads by karyotyping to prevent that balanced Robertsonian or whole arm translocations are missed in few respective cases.

Our results pave the road to a second phase that would aim at evaluating the clinical utility of the technology for all patients referred for cytogenetic investigation and assess the added-value in terms of diagnostic yield (detection of novel SVs) and genetic counselling. In fact, samples currently investigated with CNV-microarray could have undetected balanced structural variations, or other pathogenic SVs in complex regions of the genome that remain inaccessible to CNV-microarray and NGS detection, as suggested by most recent findings in singleton research cases.^47; 51^ Similarly, patients who have normal karyotype could bear variants that were undetected because they are below the karyotype resolution. Furthermore, the absence of sequencing could be preferred in some cases to avoid undesired incidental findings especially for some patients referred for reproductive disorders. The sensitivity of optical mapping to detect a repeat-contraction related disease such as Facio-Scapulo-Humeral Muscular Dystrophy (FSHD)^52; 53^ opens up new perspectives for the detection of expansion diseases such as Fragile X syndrome or Huntington disease and SVs on the Y chromosome which is rich in repetitive sequences and still challenging for sequencing.

To conclude, this is the first clinical study to validate genome-mapping as a solid alternative approach to karyotyping, FISH and CNV-microarray for the detection of chromosomal aberrations in constitutional diseases. We showed that optical mapping is capable of reaching 100% concordance, while detecting all different types of chromosomal anomalies including aneuploidies, CNVs as well as balanced chromosomal abnormalities and complex chromosomal rearrangements.

## Acknowledgments

We are thankful to the Department of Human Genetics in Nijmegen, especially Helger Yntema, Lisenka Vissers, Marcel Nelen, and Han Brunner for providing support and critical feedback. We are grateful to the Radboudumc Genome Technology Center for infrastructural and computational support. A. Hoischen, Ph.D. was supported by the Solve-RD project. The Solve-RD project has received funding from the European Union’s Horizon 2020 research and innovation program under grant agreement No 779257. SOLVE-RD. This research was part of the Netherlands X-omics Initiative and partially funded by NWO, project 184.034.019 and Radboud Institute for Molecular Life Sciences PhD grants (to A. Hoischen). TM was supported by the Sigrid Jusélius Foundation.

We are grateful to the French “Agence de la Biomédecine” and the APHP.center Paris university hospitals for their financial support the French part of the project (granted to L. El Khattabi; AOR 2018 “ART, prenatal diagnosis and genetic diagnosis” and Merri-SERI 2019, respectively). We would like also to thank Faten Hsoumi (Cytogenetics department, Cochin Hospital, Paris, France) for getting some CNV-microarray images, Emilie Chopin and Isabelle Rouvet (Cellular biotechnology center, Hospices Civils de Lyon, France) for providing lymphoblastoid cell lines, the Gentyane facility staff for providing genome imaging service for samples from Clermont-Ferrand hospital (France), and the clinical geneticists and cytogeneticists from the French ANI project whom patients were included in the present study (Bruno Delobel, Bénédicte Duban-Bedu, Dominique Martin-Coignard, Marc Planes, Céline Freihuber, Jean-Pierre Siffroi, Florence Amblard, Marlène Rio, Laurence Lohman, Véronique Paquis, Françoise Devillard, Bertrand Isidor, James Lespinasse, Gwenaël Nadeau and Laurent Pasquier). The ANI project aims at better characterizing *de novo* apparently balanced chromosomal rearrangements associated with intellectual disability using high throughput sequencing (ANI study is supported by the French Ministry of Health (DGOS) and the French National Agency for Research (ANR) PRTS 2013 grant to C. Schluth-Bolard, n° PRTSN1300001N).

We would like to acknowledge support from scientists and staff at Bionano Genomics including Alex Hastie, Andy Pang, Lucia Muraro, Kees-Jan Francoijs, Sven Bocklandt, Yannick Delpu, Mark Oldakowski, Ernest Lam, Thomas Anantharaman, Scott Way, Henry Sadowski, Amy Files, Carly Proskow.

## Web Resources

Bionano Access™: https://bionanogenomics.com/support/software-downloads/#bionanoaccess

## Abbreviations

CNV: copy number variant
DD: developmental disorder
DLE-1: direct Labeling Enzyme-1
DLS: direct Label and Stain
EDTA: ethylenediaminetetraacetic acid
FISH: fluorescence in situ hybridization
FSHD: facioscapulohumeral dystrophy
gDNA: genomic DNA
ID: intellectual disability
i.e.: id est (that is)
MCA: multiple congenital malformations
NGS: next generation sequencing
SV: structural variant
UHMW: ultra-long high molecular weight
WES: whole exome sequencing
WGS: whole genome sequencing

## Supplemental Data Description

Supplemental Data includes 6 figures and 3 tables.

**Supplementary Figure 1.** Workflow of the Bionano technique. For this study, 85 samples for whom extra material was available were included. Ultra-high molecular weight DNA was extracted using the Bionano solution phase DNA isolation method. Labeling was done using the DLE-1 chemistry. High resolution imaging of DNA molecules was done on Bionano Saphyr instruments. As different centers were included, different amount of data was produced (~800Gbp for Radboud, ~300Gbp for the French centers), and samples were analyzed using different software versions (3.4.1 and 3.5). A *de novo* assembly was performed, and both SVs and CNVs were called.

**Supplementary Figure 2.** Representative Bionano CNV profiles for different samples. A) Sample 2. Loss of 22q11.21(18645354_21465660). B) Sample 8. Gain of 17p12(14087934_15436895). C) Sample 70. Loss of 6q14.1q14.3(76385698_86884355), and gain of 6q16.1(97661978_98726638). D) Genome-wide CNV view (available in Bionano Solve v1.5) of sample 73 with E) chromosome 8 highlighted (showing a deletion) and F) chromosome 17 highlighted (showing a duplication). Blue: gains, Red: losses.

**Supplementary Figure 3:** Isochromosomes. A) Sample 77 (ish idic(15)(D15Z1+,SNRPN++,D15Z1+)). Left: Circos plot showing an abnormal CNV profile on chromosome 15. Top right: CNV-microarray data showing a gain on chr15. Bottom right: optical mapping data, showing a CNV profile that is nearly identical to the CNV-microarray profile. Numbers present fractional copy numbers. B) Sample 78 (46,X,idic(X)(p11.21)). Left: Circos plot showing a CNV baseline suggesting one copy of chromosome X (compared to the CNV line of chr22 partially shown on the left side). Additionally, the CNV profile shows a mosaic “gain” (compared to the baseline) on part of the chrX p-arm and the whole q-arm. Top right: CNV-microarray data showing a global loss on chrX (compared to a 46,XX control sample). However, the degree of loss varies within the chromosome consistent with a mosaic 45,X/46,X,idic(X)(p11.21) karyotype. Bottom right: optical mapping data showing a CNV profile that is nearly identical to the CNV-microarray profile. Numbers present fractional copy numbers. Red box shows parts of the chromsome 15 and X respectively that make up the iso-chromsomes. Grey box indicates the centromere (15 and X) and/or acrocentric p-arm (15).

**Supplementary Figure 4.** Genome imaging breakpoint detection for translocation t(9;17)(p13;q21), disrupting the gene *KANSL1* (patient 49). The two green bars represent the references of chromosomes 9 and 17, respectively. The mint bar in between represents the genome map of the translocation. The blue bar underneath represents the *KANSL1* gene. Small vertical black lines represent identified labels, and the red vertical lines indicate the translocation breakpoints, with an uncertain region of 3,828 bp in between shown in purple. The breakpoints are located between basepair-positions 35,771,617 and 35,773,383 on chromosome 9, and between 44,137,912 and 44,141,740 on chromosome 17.

**Supplementary Figure 5:** Complex sample 52. A) Karyotype of sample 52, interpreted as 46,XY,der(8)t(8;22)(q12;q12),der(13)t(8;13)(q31;q23),der(14)t(14;15)(q11.2;q25),der(15)t(14;15)(q21;q24),der(22)t(13;22)(q31.1;p11.2). B) FISH of sample 52, using FISH probes wcp8 (red), wcp14 (green). C) FISH of sample 52, using FISH probes wcp8 (green), wcp13 (red). D) FISH of sample 52, using FISH probes wcp15 (green), wcp22 (red). E) Bionano circos plot, showing different translocations t(8;13), t(8;14), t(14,15), and intrachromosomal translocations on chr 8 and chr 15.

**Supplementary Figure 6:** Complex sample 55. A) Karyotype of sample 55, interpreted as 46,X,idic(Y)(q11.22),t(5;8)(q23;q24),t(5;11)(p12;p13)[32/50]/45,X,t(5;8)(q23;q24),t(5;11)(p12;p13)[10/50]/47,XY,idic(Y)(q11.22),t(5;8)(q23;q24),t(5;11)(p12;p13)[8/50]. B) FISH of sample 55, showing the translocations t(5;8) (left) and t(5;11) (right). C) Bionano circos plot of sample 55, showing the translocations t(5;8), t(5;11) and an intrachromosomal translocations 5. D) Bionano genome maps, showing the intrachromosomal translocation on chromosome 5, which is disrupting the gene *GHR*.

Supplementary Table 1: Comparison of previous diagnostic findings with genome imaging results

Supplementary Table 2: Technical performance of genome imaging

Supplementary Table 3: Overall numbers of variants per sample

## References

1. Vissers, L.E., Veltman, J.A., van Kessel, A.G., and Brunner, H.G. (2005). Identification of disease genes by whole genome CGH arrays. Hum Mol Genet 14 Spec No. 2, R215–223.

2. Speicher, M.R., and Carter, N.P. (2005). The new cytogenetics: blurring the boundaries with molecular biology. Nat Rev Genet 6, 782–792.

3. Smeets, D.F. (2004). Historical prospective of human cytogenetics: from microscope to microarray. Clin Biochem 37, 439–446.

4. Chantot-Bastaraud, S., Ravel, C., and Siffroi, J.P. (2008). Underlying karyotype abnormalities in IVF/ICSI patients. Reprod Biomed Online 16, 514–522.

5. Hofherr, S.E., Wiktor, A.E., Kipp, B.R., Dawson, D.B., and Van Dyke, D.L. (2011). Clinical diagnostic testing for the cytogenetic and molecular causes of male infertility: the Mayo Clinic experience. J Assist Reprod Genet 28, 1091–1098.

6. De Braekeleer, M., and Dao, T.N. (1990). Cytogenetic studies in couples experiencing repeated pregnancy losses. Hum Reprod 5, 519–528.

7. De Braekeleer, M., and Dao, T.N. (1991). Cytogenetic studies in male infertility: a review. Hum Reprod 6, 245–250.

8. Miller, D.T., Adam, M.P., Aradhya, S., Biesecker, L.G., Brothman, A.R., Carter, N.P., Church, D.M., Crolla, J.A., Eichler, E.E., Epstein, C.J., et al. (2010). Consensus statement: chromosomal microarray is a first-tier clinical diagnostic test for individuals with developmental disabilities or congenital anomalies. Am J Hum Genet 86, 749–764.

9. Alkan, C., Coe, B.P., and Eichler, E.E. (2011). Genome structural variation discovery and genotyping. Nat Rev Genet 12, 363–376.

10. Schinzel, A. (2001). Catalogue of unbalanced chromosome aberrations in man.

11. van Karnebeek, C.D., Jansweijer, M.C., Leenders, A.G., Offringa, M., and Hennekam, R.C. (2005). Diagnostic investigations in individuals with mental retardation: a systematic literature review of their usefulness. Eur J Hum Genet 13, 6–25.

12. de Vries, B.B., Pfundt, R., Leisink, M., Koolen, D.A., Vissers, L.E., Janssen, I.M., Reijmersdal, S., Nillesen, W.M., Huys, E.H., Leeuw, N., et al. (2005). Diagnostic genome profiling in mental retardation. Am J Hum Genet 77, 606–616.

13. Gilissen, C., Hehir-Kwa, J.Y., Thung, D.T., van de Vorst, M., van Bon, B.W., Willemsen, M.H., Kwint, M., Janssen, I.M., Hoischen, A., Schenck, A., et al. (2014). Genome sequencing identifies major causes of severe intellectual disability. Nature 511, 344–347.

14. Lionel, A.C., Costain, G., Monfared, N., Walker, S., Reuter, M.S., Hosseini, S.M., Thiruvahindrapuram, B., Merico, D., Jobling, R., Nalpathamkalam, T., et al. (2018). Improved diagnostic yield compared with targeted gene sequencing panels suggests a role for whole-genome sequencing as a first-tier genetic test. Genet Med 20, 435–443.

15. Stavropoulos, D.J., Merico, D., Jobling, R., Bowdin, S., Monfared, N., Thiruvahindrapuram, B., Nalpathamkalam, T., Pellecchia, G., Yuen, R.K.C., Szego, M.J., et al. (2016). Whole Genome Sequencing Expands Diagnostic Utility and Improves Clinical Management in Pediatric Medicine. NPJ Genom Med 1.

16. Chaisson, M.J.P., Sanders, A.D., Zhao, X., Malhotra, A., Porubsky, D., Rausch, T., Gardner, E.J., Rodriguez, O.L., Guo, L., Collins, R.L., et al. (2019). Multi-platform discovery of haplotype-resolved structural variation in human genomes. Nat Commun 10, 1784.

17. Mantere, T., Kersten, S., and Hoischen, A. (2019). Long-Read Sequencing Emerging in Medical Genetics. Front Genet 10, 426.

18. Merker, J.D., Wenger, A.M., Sneddon, T., Grove, M., Zappala, Z., Fresard, L., Waggott, D., Utiramerur, S., Hou, Y., Smith, K.S., et al. (2018). Long-read genome sequencing identifies causal structural variation in a Mendelian disease. Genet Med 20, 159–163.

19. Mizuguchi, T., Suzuki, T., Abe, C., Umemura, A., Tokunaga, K., Kawai, Y., Nakamura, M., Nagasaki, M., Kinoshita, K., Okamura, Y., et al. (2019). A 12-kb structural variation in progressive myoclonic epilepsy was newly identified by long-read whole-genome sequencing. J Hum Genet 64, 359–368.

20. Schwartz, D.C., Li, X., Hernandez, L.I., Ramnarain, S.P., Huff, E.J., and Wang, Y.K. (1993). Ordered restriction maps of Saccharomyces cerevisiae chromosomes constructed by optical mapping. Science 262, 110–114.

21. Lam, E.T., Hastie, A., Lin, C., Ehrlich, D., Das, S.K., Austin, M.D., Deshpande, P., Cao, H., Nagarajan, N., Xiao, M., et al. (2012). Genome mapping on nanochannel arrays for structural variation analysis and sequence assembly. Nat Biotechnol 30, 771–776.

22. Chan, S., Lam, E., Saghbini, M., Bocklandt, S., Hastie, A., Cao, H., Holmlin, E., and Borodkin, M. (2018). Structural Variation Detection and Analysis Using Bionano Optical Mapping. Methods Mol Biol 1833, 193–203.

23. Wang, M., Tu, L., Yuan, D., Zhu, Shen, C., Li, J., Liu, F., Pei, L., Wang, P., Zhao, G., et al. (2019). Reference genome sequences of two cultivated allotetraploid cottons, Gossypium hirsutum and Gossypium barbadense. Nat Genet 51, 224–229.

24. Kronenberg, Z.N., Fiddes, I.T., Gordon, D., Murali, S., Cantsilieris, S., Meyerson, O.S., Underwood, J.G., Nelson, B.J., Chaisson, M.J.P., Dougherty, M.L., et al. (2018). High-resolution comparative analysis of great ape genomes. Science 360.

25. Nowoshilow, S., Schloissnig, S., Fei, J.F., Dahl, A., Pang, A.W.C., Pippel, M., Winkler, S., Hastie, A.R., Young, G., Roscito, J.G., et al. (2018). The axolotl genome and the evolution of key tissue formation regulators. Nature 554, 50–55.

26. Neveling, K., Mantere, T., Vermeulen, S., Oorsprong, M., van Beek, R., Kater-Baats, E., Pauper, M., van der Zande, G., Smeets, D., Weghuis, D.O., et al. (2020). Next generation cytogenetics: comprehensive assessment of 48 leukemia genomes by genome imaging. bioRxiv, 2020.2002.2006.935742.

27. Barseghyan, H., Delot, E.C., and Vilain, E. (2018). New technologies to uncover the molecular basis of disorders of sex development. Mol Cell Endocrinol 468, 60–69.

28. Du, C., Mark, D., Wappenschmidt, B., Bockmann, B., Pabst, B., Chan, S., Cao, H., Morlot, S., Scholz, C., Auber, B., et al. (2018). A tandem duplication of BRCA1 exons 1-19 through DHX8 exon 2 in four families with hereditary breast and ovarian cancer syndrome. Breast Cancer Res Treat 172, 561–569.

29. Levy-Sakin, M., Pastor, S., Mostovoy, Y., Li, L., Leung, A.K.Y., McCaffrey, J., Young, E., Lam, E.T., Hastie, A.R., Wong, K.H.Y., et al. (2019). Genome maps across 26 human populations reveal population-specific patterns of structural variation. Nat Commun 10, 1025.

30. Bates, S.E. (2011). Classical cytogenetics: karyotyping techniques. Methods Mol Biol 767, 177–190.

31. https://bionanogenomics.com/wp-content/uploads/2018/04/30110-Bionano-Solve-Theory-of-Operation-Structural-Variant-Calling.pdf.

32. https://bionanogenomics.com/wp-content/uploads/2018/04/30210-Introduction-to-Copy-Number-Analysis.pdf.

33. Schluth-Bolard, C., Diguet, F., Chatron, N., Rollat-Farnier, P.A., Bardel, C., Afenjar, A., Amblard, F., Amiel, J., Blesson, S., Callier, P., et al. (2019). Whole genome paired-end sequencing elucidates functional and phenotypic consequences of balanced chromosomal rearrangement in patients with developmental disorders. J Med Genet 56, 526–535.

34. Weissensteiner, M.H., Pang, A.W.C., Bunikis, I., Hoijer, I., Vinnere-Petterson, O., Suh, A., and Wolf, J.B.W. (2017). Combination of short-read, long-read, and optical mapping assemblies reveals large-scale tandem repeat arrays with population genetic implications. Genome Res 27, 697–708.

35. Redin, C., Brand, H., Collins, R.L., Kammin, T., Mitchell, E., Hodge, J.C., Hanscom, C., Pillalamarri, V., Seabra, C.M., Abbott, M.A., et al. (2017). The genomic landscape of balanced cytogenetic abnormalities associated with human congenital anomalies. Nat Genet 49, 36–45.

36. Dong, Z., Wang, H., Chen, H., Jiang, H., Yuan, J., Yang, Z., Wang, W.J., Xu, F., Guo, X., Cao, Y., et al. (2018). Identification of balanced chromosomal rearrangements previously unknown among participants in the 1000 Genomes Project: implications for interpretation of structural variation in genomes and the future of clinical cytogenetics. Genet Med 20, 697–707.

37. Kosugi, S., Momozawa, Y., Liu, X., Terao, C., Kubo, M., and Kamatani, Y. (2019). Comprehensive evaluation of structural variation detection algorithms for whole genome sequencing. Genome Biol 20, 117.

38. Monlong, J., Cossette, P., Meloche, C., Rouleau, G., Girard, S.L., and Bourque, G. (2018). Human copy number variants are enriched in regions of low mappability. Nucleic Acids Res 46, 7236–7249.

39. Luo, F. (2019). A systematic evaluation of copy number alterations detection methods on real SNP array and deep sequencing data. BMC Bioinformatics 20, 692.

40. Zhao, L., Liu, H., Yuan, X., Gao, K., and Duan, J. (2020). Comparative study of whole exome sequencing-based copy number variation detection tools. BMC Bioinformatics 21, 97.

41. Miga, K.H., Koren, S., Rhie, A., Vollger, M.R., Gershman, A., Bzikadze, A., Brooks, S., Howe, E., Porubsky, D., Logsdon, G.A., et al. (2020). Telomere-to-telomere assembly of a complete human X chromosome. Nature.

42. Lee, K.S., Choi, M., Kwon, D.W., Kim, D., Choi, J.M., Kim, A.K., Ham, Y., Han, S.B., Cho, S., and Cheon, C.K. (2020). A novel de novo heterozygous DYRK1A mutation causes complete loss of DYRK1A function and developmental delay. Sci Rep 10, 9849.

43. van Bon, B.W., Coe, B.P., Bernier, R., Green, C., Gerdts, J., Witherspoon, K., Kleefstra, T., Willemsen, M.H., Kumar, R., Bosco, P., et al. (2016). Disruptive de novo mutations of DYRK1A lead to a syndromic form of autism and ID. Mol Psychiatry 21, 126–132.

44. Ji, J., Lee, H., Argiropoulos, B., Dorrani, N., Mann, J., Martinez-Agosto, J.A., Gomez-Ospina, N., Gallant, N., Bernstein, J.A., Hudgins, L., et al. (2015). DYRK1A haploinsufficiency causes a new recognizable syndrome with microcephaly, intellectual disability, speech impairment, and distinct facies. Eur J Hum Genet 23, 1473–1481.

45. Coe, B.P., Witherspoon, K., Rosenfeld, J.A., van Bon, B.W., Vulto-van Silfhout, A.T., Bosco, P., Friend, K.L., Baker, C., Buono, S., Vissers, L.E., et al. (2014). Refining analyses of copy number variation identifies specific genes associated with developmental delay. Nat Genet 46, 1063–1071.

46. Koolen, D.A., Kramer, J.M., Neveling, K., Nillesen, W.M., Moore-Barton, H.L., Elmslie, F.V., Toutain, A., Amiel, J., Malan, V., Tsai, A.C., et al. (2012). Mutations in the chromatin modifier gene KANSL1 cause the 17q21.31 microdeletion syndrome. Nat Genet 44, 639–641.

47. Demaerel, W., Mostovoy, Y., Yilmaz, F., Vervoort, L., Pastor, S., Hestand, M.S., Swillen, A., Vergaelen, E., Geiger, E.A., Coughlin, C.R., et al. (2019). The 22q11 low copy repeats are characterized by unprecedented size and structural variability. Genome Res 29, 1389–1401.

48. Chaisson, M.J., Wilson, R.K., and Eichler, E.E. (2015). Genetic variation and the de novo assembly of human genomes. Nat Rev Genet 16, 627–640.

49. Cameron, D.L., Di Stefano, L., and Papenfuss, A.T. (2019). Comprehensive evaluation and characterisation of short read general-purpose structural variant calling software. Nat Commun 10, 3240.

50. Mahmoud, M., Gobet, N., Cruz-Davalos, D.I., Mounier, N., Dessimoz, C., and Sedlazeck, F.J. (2019). Structural variant calling: the long and the short of it. Genome Biol 20, 246.

51. Cretu Stancu, M., van Roosmalen, M.J., Renkens, I., Nieboer, M.M., Middelkamp, S., de Ligt, J., Pregno, G., Giachino, D., Mandrile, G., Espejo Valle-Inclan, J., et al. (2017). Mapping and phasing of structural variation in patient genomes using nanopore sequencing. Nat Commun 8, 1326.

52. Zheng, Y., Kong, L., Xu, H., Lu, Y., Zhao, X., Yang, Y., Yu, G., Li, P., Liang, F., Jin, H., et al. (2020). Rapid prenatal diagnosis of Facioscapulohumeral Muscular Dystrophy 1 by combined Bionano optical mapping and karyomapping. Prenat Diagn 40, 317–323.

53. Dai, Y., Li, P., Wang, Z., Liang, F., Yang, F., Fang, L., Huang, Y., Huang, S., Zhou, J., Wang, D., et al. (2020). Single-molecule optical mapping enables quantitative measurement of D4Z4 repeats in facioscapulohumeral muscular dystrophy (FSHD). J Med Genet 57, 109–120.

