## Supplemental Figures 1-6 for "Next generation cytogenetics: genome-imaging enables comprehensive structural variant detection for 100 constitutional chromosomal aberrations in 85 samples"

**Supplementary Figure 1:** Workflow of the Bionano technique.

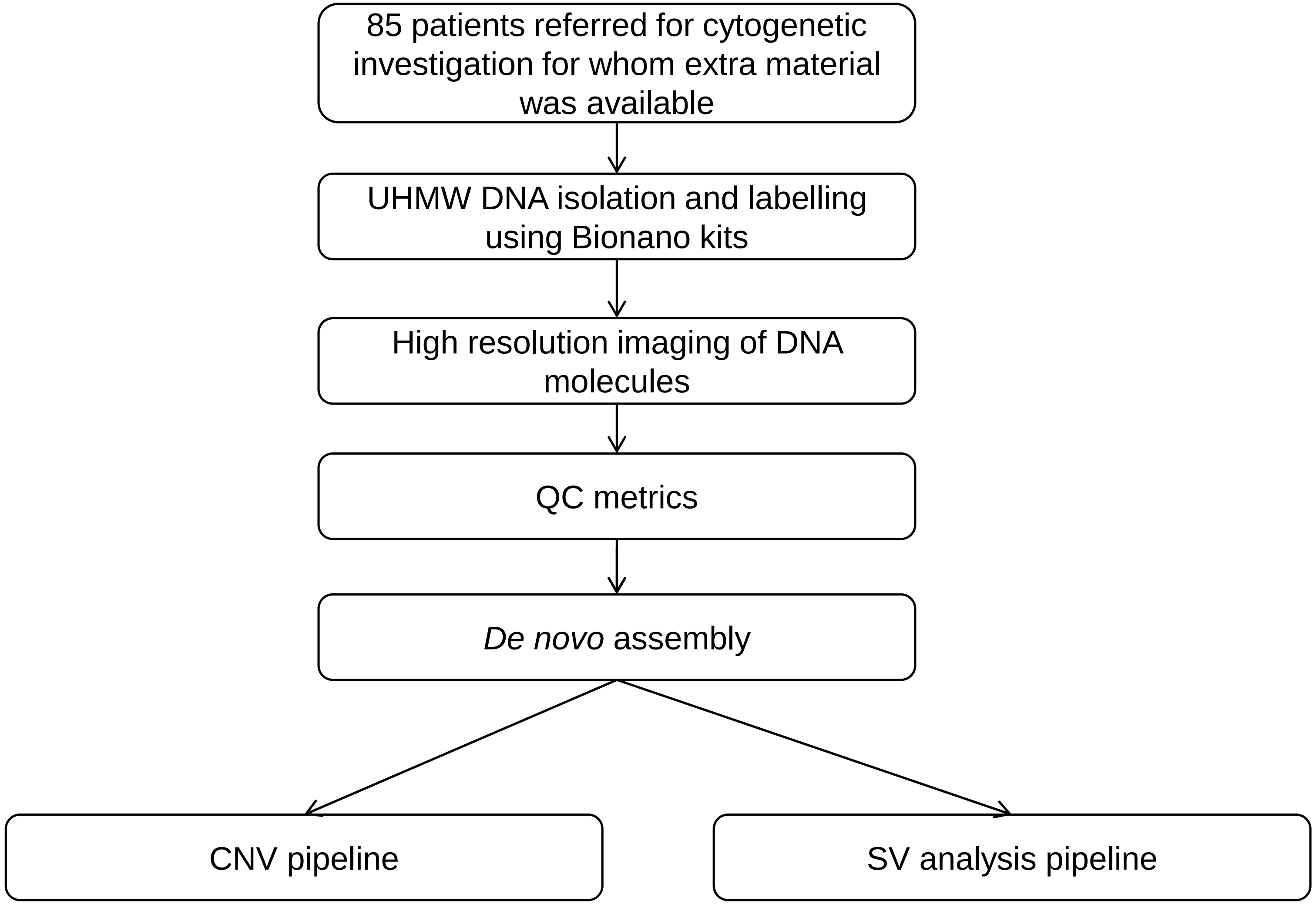

**Supplementary Figure 2:** Representative Bionano CNV profiles for different samples

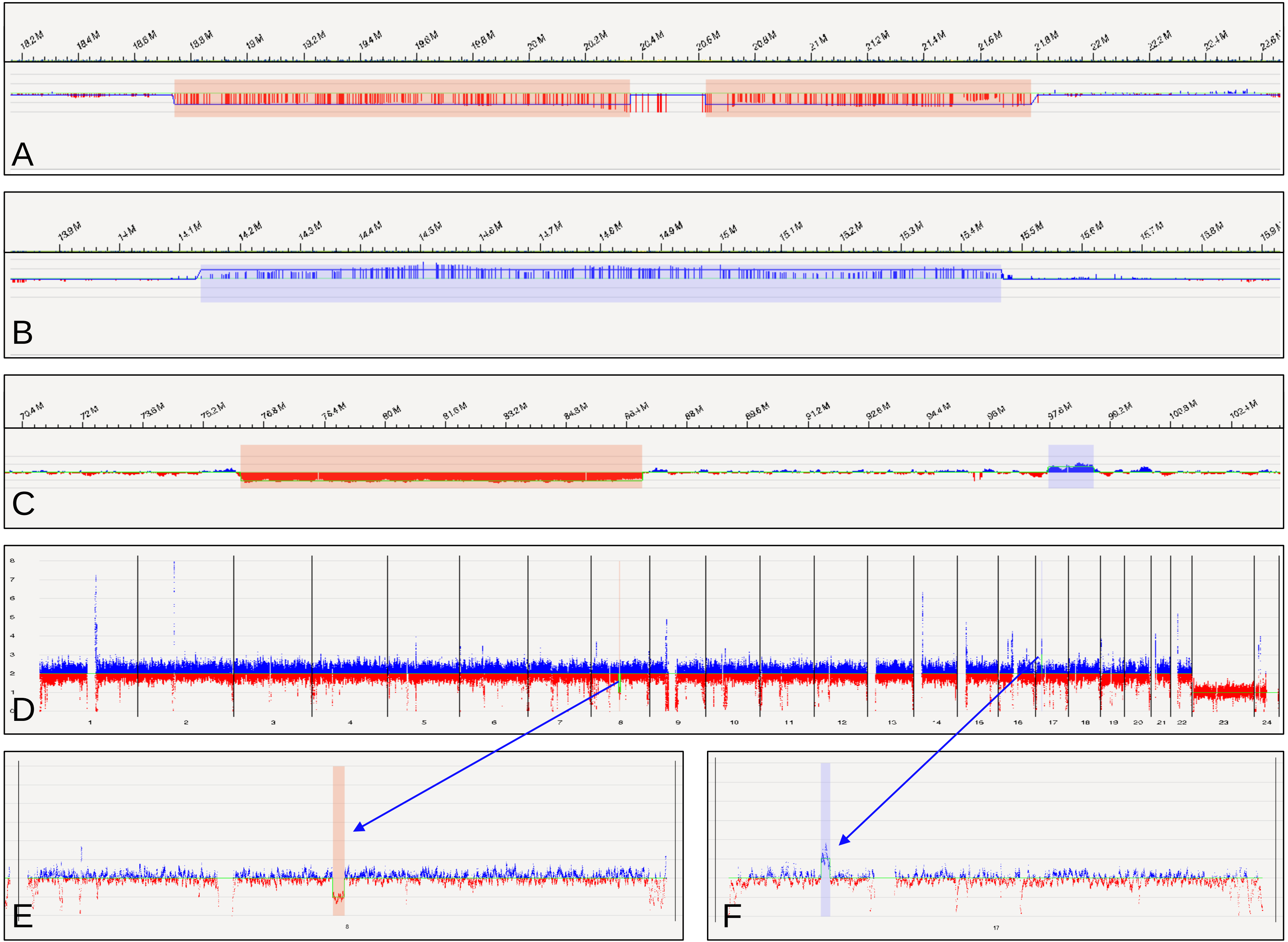

**Supplementary Figure 3**: Isochromosomes

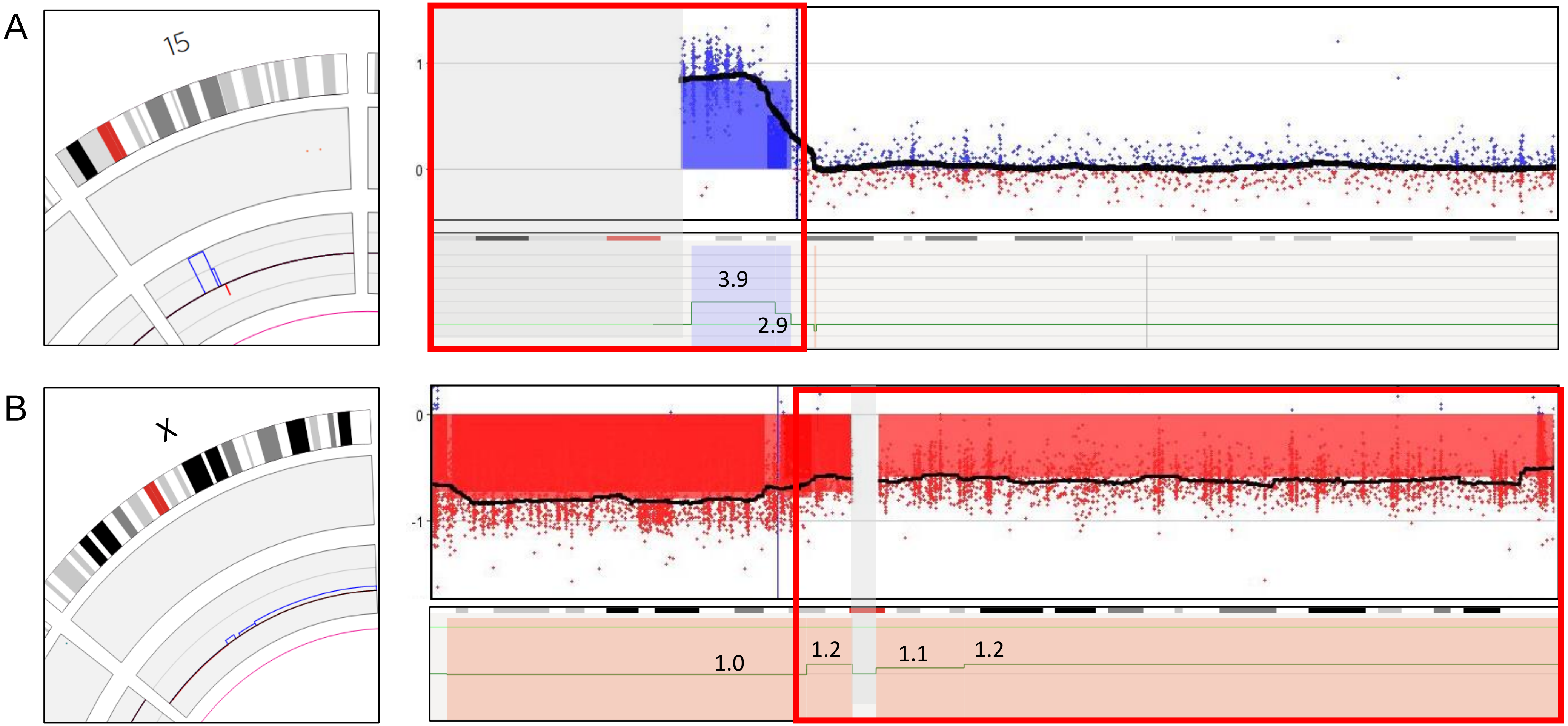

**Supplementary Figure 4:** Genome imaging breakpoint detection for translocation t(9;17)(p13;q21), disrupting the gene *KANSL1* (patient 49)

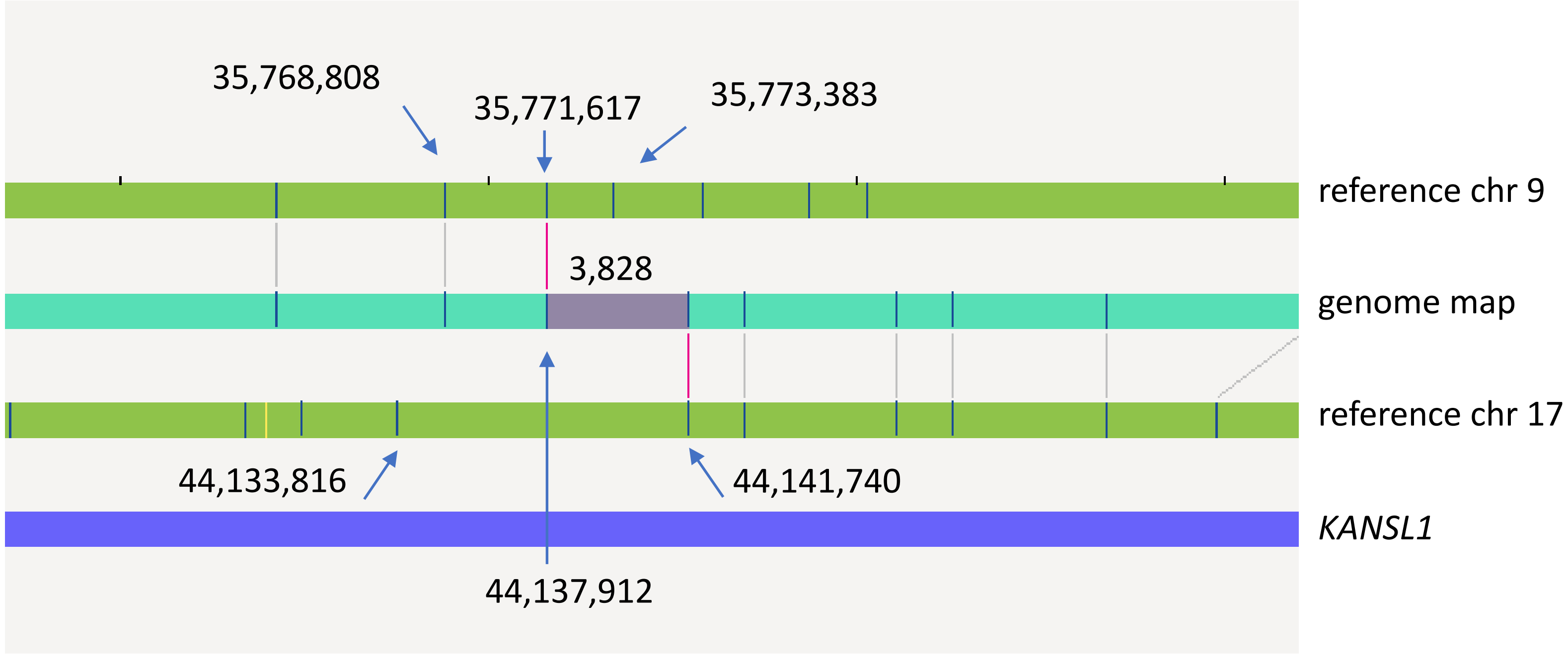

**Supplementary Figure 5**: Complex sample 52

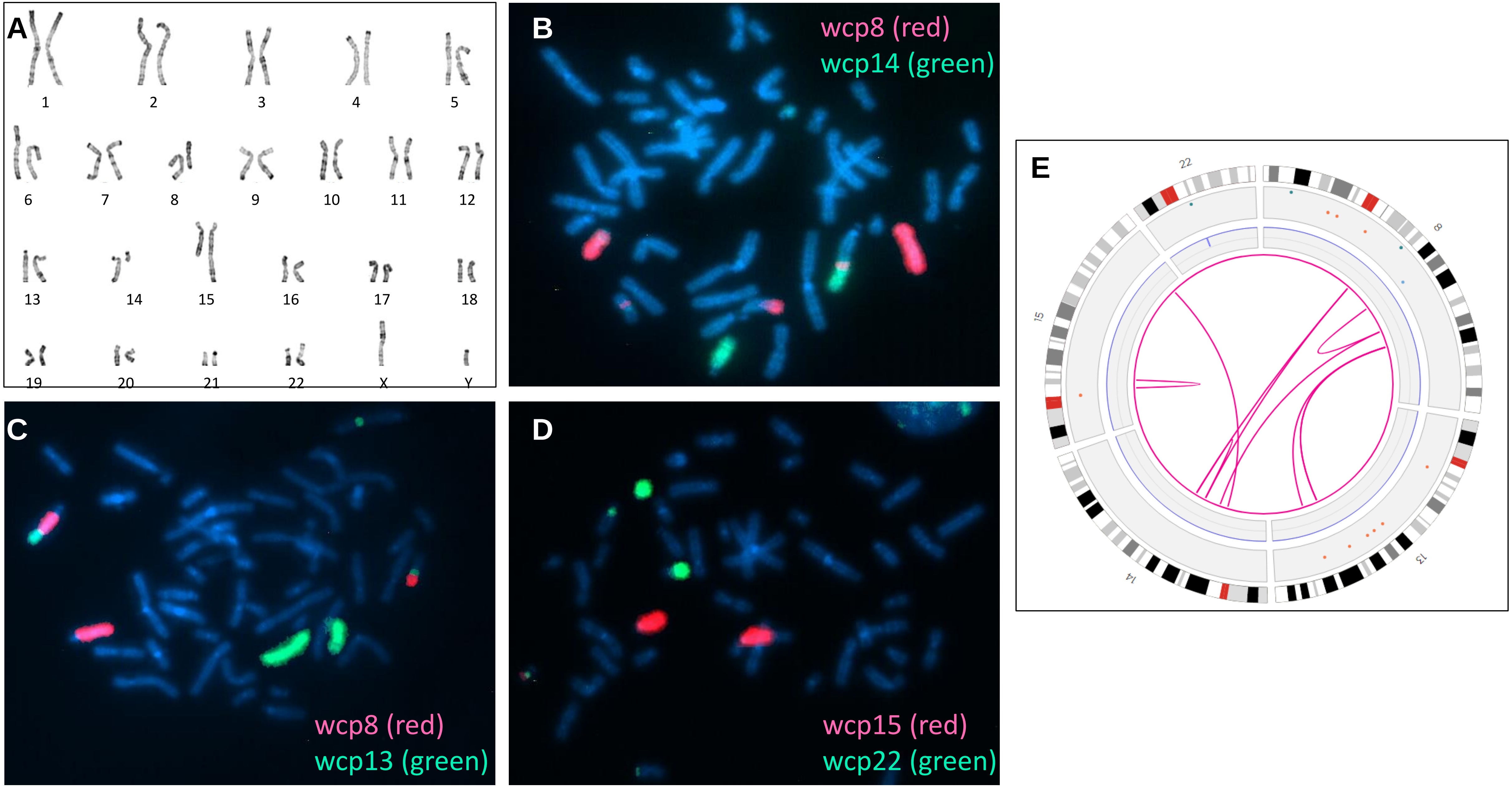

**Supplementary Figure 6**: Complex sample 55

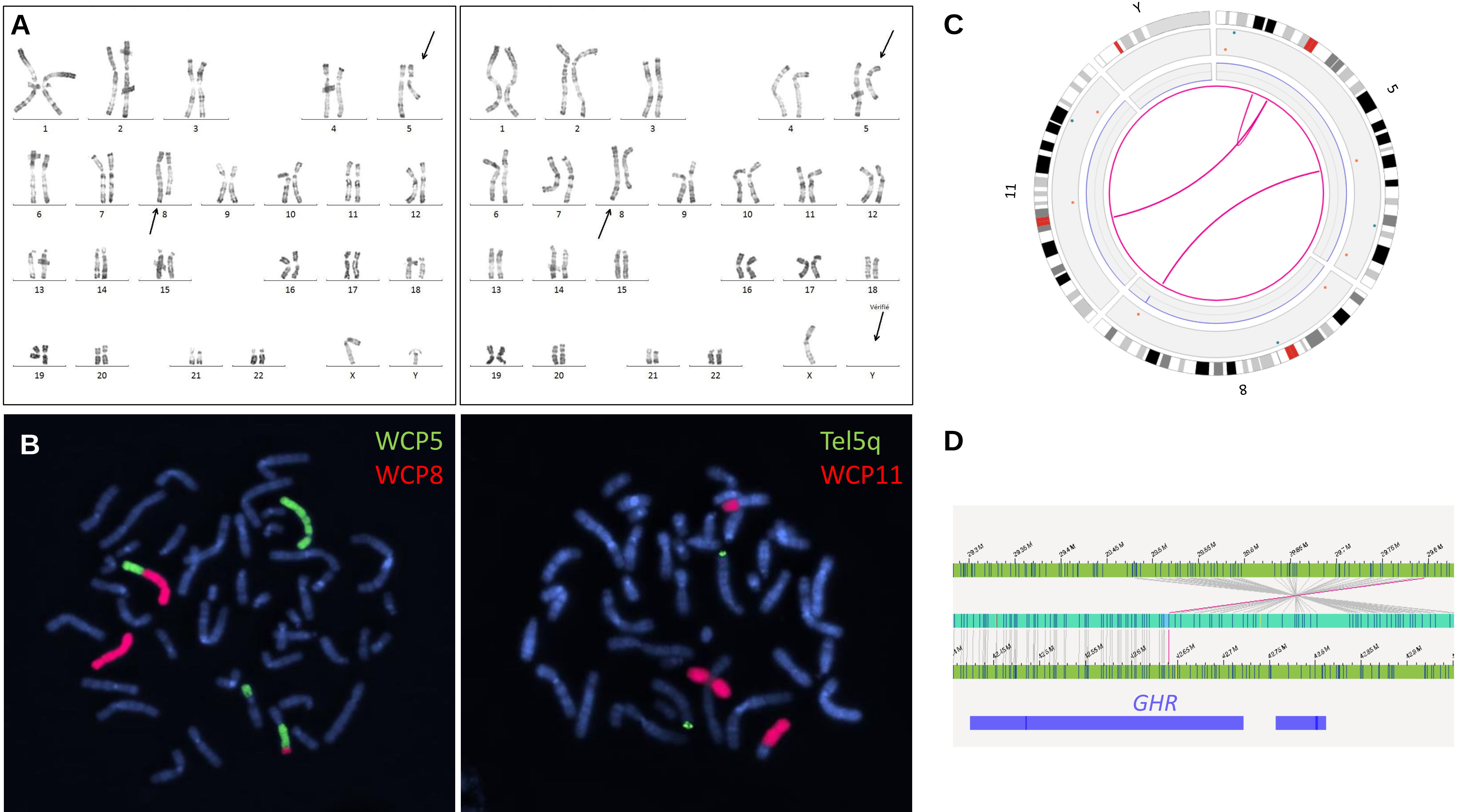
